# Effects of sex and gonadal hormones on manually segmented hypothalamic and pituitary gland volumes in young healthy adults

**DOI:** 10.1101/2023.07.17.549333

**Authors:** Sherri Lee Jones, Chloe Anastassiadis, Matthieu Dupuis, Jens Pruessner

## Abstract

The hypothalamus and pituitary regulate, amongst other functions, third order endocrine systems, and their volumes have been associated with normal and pathological outcomes. Yet, there are very few studies that examine their combined structural variations *in vivo*. This is due, in part, to their small size and a lack of comprehensive image segmentation protocols. In the current project we acquired high-resolution T1- (1mm isotropic) and T2-weighted (0.4mm in plane resolution) 3T magnetic resonance images (MRI) of the hypothalamus and pituitary gland, as well as salivary estradiol and testosterone from 31 (17M, 14F) young healthy adults. Women reported oral contraceptive use. Image preprocessing included non-uniformity correction, signal intensity normalization and standard stereotaxic space registration. We applied a comprehensive manual segmentation protocol of the whole hypothalamus, with detailed segmentation of the pituitary stalk, the anterior and posterior pituitary gland, and the posterior bright spot. We also propose a novel medial-lateral hypothalamic parcellation into medial preoptic, periventricular (PVN), and lateral hypothalamic regions. The protocol yielded good inter- (range: 0.78-0.92) and intra-rater (range: 0.79-0.94) Dice kappa overlap coefficients. We detected sex differences of the whole hypothalamus and each hemisphere, and a trend for the right preoptic region to be larger in males than in females, with a moderate effect size. Sex differences were maintained or enhanced when covarying for estradiol, but not when covarying for testosterone. In addition, testosterone was associated with the volume of the PVN, but only in women. In summary, these results suggest that there are morphometric differences at the level of the pituitary and hypothalamus that are likely driven by central regulation of gonadal hormones. The here described protocol allows the structural investigation of neuroendocrine effects in the central nervous system.

## Introduction

The hypothalamus is a sexually dimorphic neural structure and through its connections to the pituitary gland, is the body’s master neuroendocrine regulator. The hypothalamus is primarily involved in regulating autonomic functions, bodily homeostasis, and emotional responses. It has multiple afferent and efferent connections, including bi-directional connections to the cerebral cortex, hippocampal formation, amygdala and brainstem (Saper et al., 2004; Saper & Lowell, 2014), and controls third-order feedback endocrine systems through signals to the pituitary gland (also called the hypophysis). Despite its small size, its structurally and functionally distinct nuclei serve critically important functions in the organism. For example, the medial preoptic nucleus, arcuate nucleus, and paraventricular nucleus send trophic factors via the hypophyseal portal system to stimulate releasing hormones from the anterior pituitary gland (adenohypophysis) to control regenerative, metabolic and reproductive function. Cell bodies located in the paraventricular nucleus and supraoptic nucleus project to and comprise the nerve tissue of the posterior pituitary gland (neurohypophysis), to release oxytocin and vasopressin into the blood stream for regulation of social behaviors (Amar & Weiss, 2003; Saper et al., 2004; Saper & Lowell, 2014). Recently, post-mortem and *in-vivo* MRI studies have shown hypothalamic volume to be affected in mood disorders (Bao et al., 2005; Manaye et al., 2005; S. Schindler et al., 2019; Stephanie Schindler et al., 2012), psychotic disorders (Klomp et al., 2012) Parkinson’s (Breen et al., 2016), frontotemporal dementia (Bocchetta et al., 2015), and Alzheimer’s disease (Callen et al., 2001). Despite the importance of the hypothalamus and pituitary in neuroendocrinology, there exists no single comprehensive protocol to study the morphology of the hypothalamus and pituitary, and their substructures, *in vivo*, and no protocol as far as we know, that also accounts for circulating sex steroid hormones.

The last decade has brought important advances in delineating and parceling the hypothalamus on standard MRI images (Bocchetta et al., 2015; Goldstein et al., 2007; Makris et al., 2013; Piguet et al., 2010; Stephanie Schindler et al., 2013). Those that have parceled the hypothalamus have done so into three anteroposterior regions: the anterior hypothalamus, tuberal region, and posterior hypothalamus, and further differentiate the anterior and tuberal regions into superior and inferior parcels (Billot et al., 2020; Makris et al., 2013; Stephanie Schindler et al., 2013), with each parcel containing specific nuclei (Makris et al., 2013). Importantly, none of the existing hypothalamic segmentation protocols, have defined the medial-lateral divisions.

Despite advances in hypothalamic segmentation, some important limitations remain. First, most protocols exclude portions of the anterior preoptic area (POA), a region implicated in sex-specific behavior and physiology, containing sexually-dimorphic nuclei including the classic sexually dimorphic nucleus (SDN-POA) (Bao & Swaab, 2011; Gorski et al., 1978; Swaab, 1995; Swaab & Fliers, 1985). This nucleus is also known as the first interstitial nucleus of the anterior hypothalamus (Allen & Gorski, 1990), or the intermediate nucleus (H Braak & Braak, 1987; Yuri Koutcherov et al., 2007). The importance of the POA has been shown in a number of sexually differentiated motivated behaviors including maternal behavior (Champagne et al., 2003, 2006; Numan & Stolzenberg, 2009), sexual behaviors (Graham et al., 2015; Graham & Pfaus, 2013; Matuszewich et al., 2000; Paredes, 2003; Whitney, 1986), and gender identification (Garcia-Falgueras & Swaab, 2008; Hoekzema et al., 2015). Thus, being able to fully include the POA would improve our understanding of sex differences in the brain and the behaviors this region subserves.

Second, the ventral boundary of the hypothalamus at the level of the pituitary stalk has not been clearly defined. This may not be particularly problematic when segmented on standard T1 isotropic images due to it comprising only a small area of the ventral diencephalon. However, morphological features of the stalk are clinically important in neoplastic, inflammatory, and infectious disorders (Satogami et al., 2010), and an objective definition is required for any protocol aiming to segment the pituitary gland in combination with the hypothalamus, given that the stalk is the critical link between the two structures.

Third, existing protocols exclude portions of the dorsal boundary of the paraventricular nucleus (PVN) at the level of the fornix. The PVN is one of the most important hypothalamic nuclei involved in stress system regulation and likely plays a key role in stress-related psychopathologies. Moreover, the PVN is an important integrative nucleus for the interaction between the hypothalamic-pituitary adrenal (HPA) and gonadal (HPG) axes (Toufexis et al., 2014). Thus, it might be particularly relevant for understanding stress-dependent psychopathologies that differ between the sexes (Viau, 2002).

Fourth, the hypothalamus can be divided into lateral, medial and periventricular regions (Saper et al., 2004; Schönknecht et al., 2013), which is lacking from existing protocols that subdivide it along the rostral-caudal plane. This is due in part to the low resolution of standard scans relative to its cytoarchitecture, as well as intermingled white and gray matter within the structure making it particularly difficult to discriminate the lateral boundaries on standard T1 weighted images. Yet a medial-lateral parcellation will allow a focus on the periventricular region that contains the PVN, an anterior medial region containing the POA, as well as a parcellation of the lateral hypothalamus, widely known to be important in appetite regulation and food intake, including pathological obesity (Saper et al., 2004; Saper & Lowell, 2014).

Discrimination of the two structurally and functionally distinct pituitary lobes on standard structural MRI images is poorly defined. The anterior lobe (adenohypophysis) is comprised of at least six endocrine cells types, while the posterior lobe (neurohypophysis) is comprised of axon terminals from neural cell bodies that originate in the supraoptic and paraventricular nuclei from which vasopressin and oxytocin are produced (Castillo, 2005). While some have examined the anterior and posterior glands separately (Takano et al., 1999), typically the posterior bright spot, a hyperintense signal on T1-weighted images, is considered as the posterior pituitary gland and to represent its functional integrity (Côté et al., 2014; Kilday et al., 2014; Yamamoto et al., 2012). Using the bright spot as a volumetric measure of the posterior pituitary is problematic because there is controversy about what the bright spot represents. The majority of the current literature suggests that the bright spot represents vasopressin containing cells (Kitagawa et al., 2017; Lee et al., 2001; Sato et al., 1995), and therefore may reflect an individual’s state of hydration (Castillo, 2005). Thus, it seems unreasonable to consider the bright spot as the entire posterior pituitary gland based on its signal intensity, because a lack of bright spot (i.e., hypersecretion of vasopressin) would then imply an absent posterior pituitary gland. Others have defined the posterior pituitary as the part of the pituitary gland lying between the posterior aspect of the pituitary stalk and the anterior margin of the dorsum sellae (Yamamoto et al., 2012), however the authors did not provide volumetric analysis. Moreover, although this could be a reliable method, it may ignore individual differences in pituitary gland structure.

Thus, the goal of the current study was to develop a comprehensive manual segmentation protocol of the hypothalamus and pituitary while considering salivary sex hormone levels. Specifically, the novelty of this protocol is to more comprehensively define the hypothalamic borders, to parcel it in its medial-lateral division, to include segmentation of the pituitary stalk, and to distinguish between the anterior and posterior pituitary glands on T1- weighted structural MRI scans. Because the hypothalamus and pituitary have previously been shown to be too small to be reliably segmented with standard 1mm isotropic images, we aimed to increase resolution. Images were acquired on a 3T scanner and the 1mm isotropic T1 resolution was upsampled to 0.5 x 0.5 x 1 mm. In addition, we also used high resolution T2- weighted images (0.4 mm x 0.4 mm in plane) to cross-validate the results. The inclusion of T2 weighted images further allowed exclusion of major white matter tracts such as the fornix, optic tract, and anterior commissure, as also described by Bocchetta et al., (2015). Specific attention was paid to discriminate gray and white matter in the hypothalamus, and to take into account the strong signal intensity variation between the anterior and posterior pituitary. While paying particular attention to these methodological issues in the current protocol, we aimed to objectively define “gray matter” boundaries in the hypothalamus, by defining and applying a threshold based on the individual subject’s T1 weighted, and intensity normalized, scan. Within the pituitary, we further aimed to objectively define the posterior “bright spot”, and include a manual delineation of a “dividing line” using the hypo intensity signal that lies between the anterior and posterior regions of the pituitary gland.

## Materials and Methods

### Participants

Participants were 31 (17 males, 14 females) French-mother-tongue adults with a mean age of 18.9 years (± 0.23). Participants were initially recruited at age 11 ½ from high schools located in the Monteregie region of Montreal, as a community control cohort in a prospective longitudinal study of prenatal maternal stress (Charil et al., 2010; King et al., 2012), but the protocol described here used their scans acquired at approximately 19 years of age.

Participants who were born preterm (<37 weeks) and mothers that reported major negative life events (e.g., death of relative, serious illness of self/relative) (Sarason et al., 1978) during pregnancy were excluded.

The work described has been carried out in accordance the Declaration of Helsinki for experiments involving humans. Procedures were approved by the Research Ethics Board of the Douglas Hospital Research Center. All participants provided signed informed consent prior to their participation.

### MR Image Acquisition

T1 and T2 weighted images were obtained on a 3T Siemens Magnetom scanner using a 12-channel head coil. T1-weighted MP-RAGE (magnetization-prepared rapid gradient-echo) images were acquired in the sagittal plane with the following parameters: TR/TE=2400/2.43ms, flip angle 8 degrees; 192 slices; resolution 256, FoV=256; bandwidth 183 Hz/Px, 1mm isotropic and SINC interpolated to 0.5 x 0.5 x 1mm. T1 scan time was 10m16s per subject. T2-weighted images were acquired using a Turbo Spin Echo (TSE) sequence, in the anterior-posterior plane, TR/TE: 9630/85ms, flip angle 150 degrees, resolution 0.4 x 0.4 x 1mm, FoV: 150mm, 36 slices, bandwidth 107 Hz/Px, echo spacing 17.1ms). T2 scan time was 8m32s per subject.

### Hormone Contraceptive Use and Hormone Assays

Ten of the 14 women were taking hormonal contraceptives. Passive drool saliva samples for estradiol and testosterone were acquired within an hour of the scan (mean time of day=13h 28m, SD=2:58; collection time varied between 9 am and 7 pm). Salivary 17-β-estradiol and testosterone levels were assayed using Salimetrix (Pennsylvania, USA) enzyme-linked immunoassay kits (kits 1-3702 and 1-2402). All samples were run in duplicate and averaged. Salivary 17-β-estradiol levels that fell below the kit detection limit <1.0pg/mL, were assigned a value of 1.0pg/mL for statistical analysis.

### MR Image Analysis

Total brain volume including cerebrospinal fluid (TBV+CSF) and excluding (TBV-CSF) cerebral spinal fluid (CSF) were measured using the Brain Extraction based on nonlocal Segmentation Techniques (BEaST) method, which is based on nonlocal segmentation in a multi- resolution framework, and excludes the skull, skin, fat, muscles, dura, eyes, bone, exterior blood vessels and exterior nerves (Eskildsen et al., 2012), and performed using the open source pipeline BPIPE (https://github.com/CobraLab/minc-bpipe-library). All images were carefully examined to ensure proper segmentation of brain tissue. All scans passed this quality control.

All images were preprocessed using the open-source minc-toolkit software suite developed at the Montreal Neurological Institute (http://bic-mni.github.io) prior to segmentation. First, for each subject, the T2-weighted image was rigidly aligned to the T1-weighted image using *mritoself*. Next, the T1 and aligned T2-weighted images were corrected for magnetic field non-uniformities (Sled et al., 1998), linearly registered to standard stereotaxic space, resampled onto a 0.5mm voxel grid (ICBM152b), and intensity normalized (Collins et al., 1994). Linear registration into standard stereotaxic space corrects for interindividual differences in whole brain volume (Lord et al., 2010).

Images were manually segmented using the software package DISPLAY 2.0 developed at the Brain Imaging Center of the Montreal Neurological Institute (www.bic.mni.mcgill.ca/software/Display/Display.html). This software allows three-dimensional navigation, and built-in rulers to quickly measure distances of anatomical boundaries which facilitates anatomical comparisons across subjects (e.g., width of anterior commissure in coronal plane; distance from medial edge of optic tract to 3^rd^ ventricle in coronal plane). In addition, the software allows copying a label to the subsequent brain slice while applying a given threshold, which allows for more rapid segmentation. Lastly, multiple volumes can be loaded at once, allowing multimodal image comparisons (e.g., of T1 and T2), and cross- validation of anatomical structures.

### Development of Hypothalamic Segmentation Guidelines

Hypothalamic anatomy and cytoarchitecture were derived from available histological publications (H Braak & Braak, 1987; Heiko Braak & Braak, 1992; Hofman & Swaab, 1989; Y Koutcherov et al., 2000; Yuri Koutcherov et al., 2002, 2007; Young & Stanton, 1994). The human brain atlas by Mai, Majtanik and Paxinos (Mai et al., 2015) provided an additional resource, and finally ‘BigBrain’, a digitally reconstructed post-mortem human brain at 20μm isotropic resolution (Amunts et al., 2013) was also used to cross-check and validate boundaries of the target structures.

The protocol guidelines were developed by referring to an MRI atlas (Baroncini et al., 2012) and MRI connectivity maps (Lemaire et al., 2011, 2013; Schönknecht et al., 2013) of the human hypothalamus, with reference to established reliable MRI segmentation methods (Bocchetta et al., 2015; Goldstein et al., 2007; Makris et al., 2013; Piguet et al., 2010; Stephanie Schindler et al., 2013). It is however fair to say that our protocol builds on the comprehensive guidelines introduced by Bocchetta et al. (Bocchetta et al., 2015) which refined previous manual segmentation protocols by applying information acquired from T1 and T2 weighted images (1.1mm isotropic), yet our T2 scans were acquired at a higher resolution (0.4x0.4 in plane).

### Defining the image contrast intensity threshold for segmentation

The protocol was designed for segmentation on the coronal view of the T1-weighted image using an upper and lower threshold, while cross-validating with the T2 images. A document template is provided in **Supplemental Materials Document 1 - HypPit_Thresholds_CleanDataSheet.xlsx**.

One aim of this protocol was to determine an objective definition of the voxel intensity range for labeling an individual voxel as hypothalamic/pituitary tissue. This first step was deemed important particularly for the hypothalamus which is bordered by extreme hypo- and hyper-intensity signals, due to the surrounding third ventricle and base of the diencephalon, as well as major white matter tracts that are used as standard anatomical landmarks and borders of the hypothalamus (Bocchetta et al., 2015; Goldstein et al., 2007; Makris et al., 2013; Stephanie Schindler et al., 2013), and therefore may be susceptible to rater bias in segmentation of partial volume. Thus, we define a range of voxel intensity values (i.e., an upper and lower threshold) in the segmentation. Without threshold, manual segmentation of the hypothalamus and pituitary with their subdivisions on the 0.5mm isotropic images proved to be impractical (∼5-8 hours per subject). By defining a range and applying the threshold, segmentation was facilitated, reducing segmentation time to approximately 2h per subject.

Thus, this protocol first defined the contrast range from intensity normalized images based on anatomical landmarks (3^rd^ ventricle with hypointense cerebral spinal fluid, and large white matter tracts including the optic tract, anterior commissure, and internal capsule which appear hyperintense on T1-weighted images). Next, the lower and upper limit of the contrast range was calculated to determine the gray matter/CSF and gray matter/white matter threshold, respectively (see section *Manual Tracing Protocol – Hypothalamus – Setting the contrast* below). The T2-weighted image is particularly useful for excluding white matter tracts such as the fornix, anterior commissure, optic tract, and mammillothalamic tract. The contrast range was determined following the manual selection of voxels as gray matter on the T1 images, and cross-validating on the T2-weighted image. The minimum and maximum contrast values from 8 randomly selected subject scans, were then converted to proportions of their respective contrast range, then averaged. Thus, by objectively defining the gray matter threshold on the intensity normalized images, we aimed to minimize inter and intra-rater variability in voxel selection as gray matter, white matter, or partial volume. A similar procedure was employed for the pituitary gland to determine anterior pituitary tissue from the sphenoid space, and to determine the hyperintensity of the signal to be labelled as the “posterior bright spot”. Given that the proportion for the lower contrast setting (i.e., between gray matter and cerebral spinal fluid, and between pituitary tissue and sphenoid space) was similar between raters when segmenting the hypothalamus or pituitary, the protocol set a single “lower limit” value for how hyperintense the signal should be relative to cerebral spinal fluid/sphenoid space, for segmentation of the hypothalamus, pituitary stalk, and pituitary tissue (see procedures on *Setting the image contrast* in the relevant anatomical sections below.

### Manual Segmentation Protocol

#### Setting the image contrast

##### Hypothalamus

To set the contrast, we navigated the T1 image from anterior to posterior in the coronal view to the inter-ventricular foramen. Next, the image was zoomed such that the internal capsule, anterior commissure and optic tract were visible within the window, then the contrast setting was adjusted such that each voxel showed variation in contrast. The gray matter to cerebral spinal fluid (GM-CSF) boundary was set before the gray matter to white matter (GM-WM) boundary. Setting the correct contrast was facilitated by switching to spectral color-coding view (available in Display). The segmentation threshold was then defined at 40% and 75% of the image contrast range, as described in the section

##### Development of the manual segmentation protocol

On the T2 weighted image, we then navigated 1.5mm posterior and adjusted the contrast for GM-CSF (lower setting). Next, we navigated to the anterior hypothalamus where the optic tract connects to the anterior hypothalamus and adjusted the contrast so each voxel in the anterior commissure and optic chiasm showed variation in contrast.

##### Pituitary

The pituitary gland is bordered anteriorly and ventrally by the sphenoid sinus, laterally by the cavernous sinuses, posteriorly by the dorsum sellae, and dorsally by the diaphragma sellae (as described by (Klomp et al., 2012)) as well as the pituitary stalk as described above. The GM-CSF limit determined for the hypothalamus was also applied for the pituitary. To adjust the intensity within the pituitary gland, we navigated to the midline of the brain, then in sagittal view, we located the section where the pituitary stalk clearly connected the hypothalamus to the pituitary gland, and where the anterior pituitary gland and posterior bright spot were clearly visible. Next, in sagittal view, we navigated medial-laterally the width of the stalk to the section with the largest bright spot. Then we adjusted the contrast from hyperintense towards a hypointense signal until all voxels within the bright spot showed maximum variation in contrast.

On the T2-weighted image, we navigated to the midsagittal section and adjusted the contrast such that the voxels within the sphenoid space and diaphragm sellae each had maximum variation in contrast.

#### Tracing and segmentation of the hypothalamus and pituitary gland

##### Hypothalamus

The manual segmentation protocol of the hypothalamus is summarized in **Table 1**. The current protocol extends and complements the protocol proposed by Bocchetta et al. (Bocchetta et al., 2015) that was developed for 3T MRI T1-weighted and T2-weighted images of the whole hypothalamus, which itself was modified from Schindler’s (Stephanie Schindler et al., 2013) protocol for T1 weighted images. Moreover, Bocchetta’s parcellation rules which were previously proposed by Makris et al. (Makris et al., 2013), segments five subunits of the hypothalamus into the inferior and superior regions of the anterior and tuberal region, as well as a posterior parcel. No protocol to our knowledge parcels the hypothalamus in its medial-lateral extent. Our protocol parcels the hypothalamus in its medial-lateral extent, into periventricular, medial and lateral regions (Saper et al., 2004; Schönknecht et al., 2013).

**Table 1.**
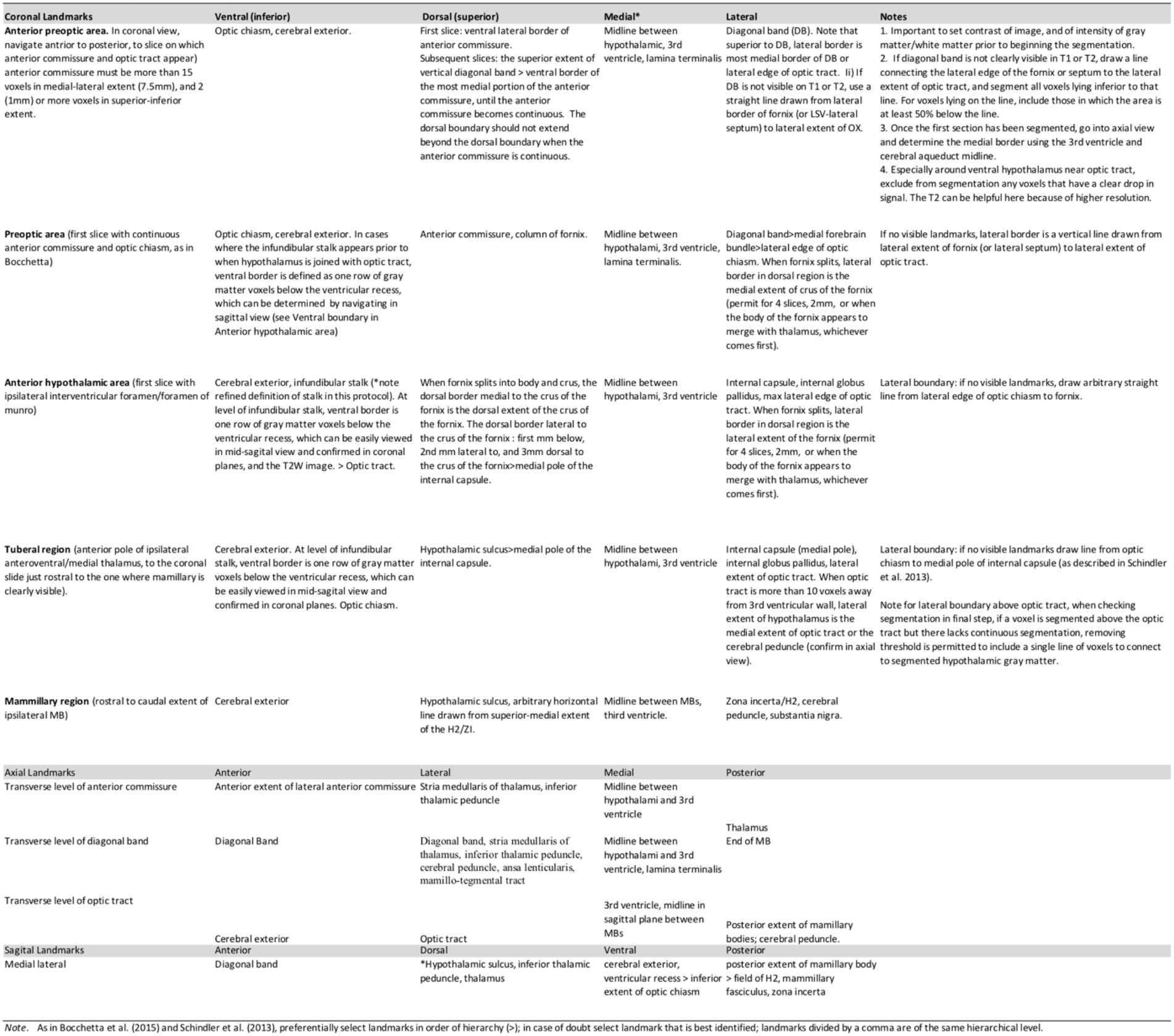
Hypothalamic segmentation guidelines.

Thus, the current protocol extends previous protocols by defining parcels for the preoptic region, the lateral hypothalamic area, as well as the periventricular region, which was possible due to higher image resolution. As such, our protocol attempts improvements to existing protocols by increasing precision in areas of interest.

A first aim was to include anterior hypothalamic tissue in the preoptic area. The first section of the anterior hypothalamus is typically considered the first slice with ipsilateral continuous anterior commissure, and in which the optic tract is attached to the brain in coronal view (Bocchetta et al., 2015; Stephanie Schindler et al., 2013). Although this is a reliable definition, it excludes portions of the anterior preoptic region, especially when segmenting on 1mm isotropic sections. With the available higher resolution images (T2’s with a 0.4mm x 0.4mm resolution in coronal plane), we defined the most rostral slice as the one containing the anterior commissure and optic tract, in which the anterior commissure is at least 7.5mm in the medial-lateral extent and at least 2 or more voxels in the superior-inferior extent (**Figure 1**).

**Figure 1.**
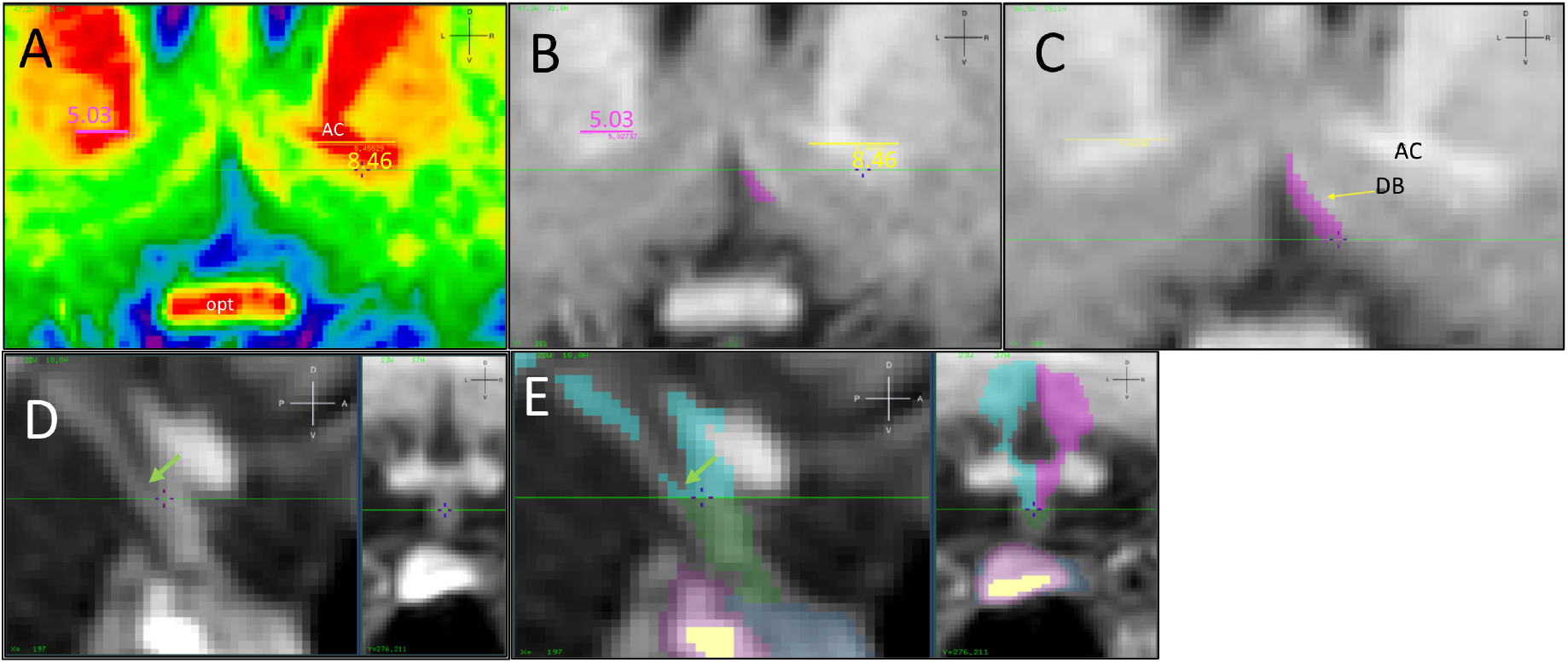
Segmenting the first section of the anterior hypothalamus. (A-C) and of the ventral hypothalamus at the level of the pituitary stalk (D-E). As in Panel A, right hemisphere, the most rostral slice is defined as the one containing the anterior commissure and optic tract, in which the anterior commissure is at least 7.5mm in the medial-lateral extent and at least 2 or more voxels in the superior-inferior extent. The AC does not meet the length definition on the left hemisphere. Determining the limits of the AC is facilitated using spectral view (Panel A) as only dark red voxels are considered, which is more difficult to define in grayscale (Panel B). Panel C shows the final segmentation of the first slide on the right hemisphere. Segmentation of ventral hypothalamus at level of pituitary stalk. Locate the voxel at the ventral extent of the infundibular recess (VXinf, green arrow in D and E), and draw a line one row of voxels below it (D). The line determines the ventral border of the hypothalamus segmented in blue, and dorsal border of the pituitary stalk which is segmented in green (E). AC: Anterior commissure; opt: optic tract. Orientation is shown in upper right corner of each panel. A: Anterior; D: dorsal; L: left; P: posterior; R: right; V: ventral. Each voxel = 0.5mm.

This definition was defined by careful morphological study of the Mai, Majtanik and Paxinos (Mai et al., 2015) atlas, and is similar to that applied by Makris et al. (Makris et al., 2013), who began segmentation on the section containing the rostral most tip of the anterior commissure and optic chiasm. We refer to this region as the anterior preoptic area (row 1 of **Table 1**). The dorsal border on the first slice was defined as the most ventral lateral border of the anterior commissure, and for subsequent slices, the most ventral medial extent of the anterior commissure. However, the dorsal border never exceeded the most superior voxels at the level of continuous anterior commissure. The lateral border in this anterior preoptic region was defined as the diagonal band (using T2 for confirmation) and at maximum the lateral edge of the optic tract. If the diagonal band was not visible, the lateral border was an arbitrary line drawn from the lateral edge of the fornix (or lateral septum if anterior to the fornix) to the lateral extent of the optic tract. The medial and ventral borders were defined as the 3^rd^ ventricle and base of the diencephalon, respectively. The sagittal orientation was verified to ensure that only hypothalamic tissue was included, and not cortical matter. Starting with the first slice with continuous anterior commissure and optic chiasm, we applied the segmentation rules of the Bocchetta (Bocchetta et al., 2015) protocol (summarized in **Table 1)**.

The ventral border of the hypothalamus at the level of the pituitary stalk has not, as far as we know, been objectively defined in any hypothalamus segmentation protocol. Because we took a neuroendocrine approach to the segmentation of the hypothalamus, we objectively defined a ventral border that would ensure the inclusion of ventrally located cell bodies (e.g., of the arcuate nucleus) in the segmentation, and that complemented existing MRI methodologies (on a 3T Siemens scanner using a standard T1-weighted MP-RAGE image sequence) in the study of the pituitary stalk (Satogami et al., 2010). The pituitary stalk was identified using the sagittal plane, and by navigating from the medial to lateral extent of the stalk we were able to identify the ventral tip of the infundibular recess of the third ventricle, which was defined as the inferior-most voxel (VXinf) within the ventricle that was showing a hypointense signal with reference to its surrounding voxels. The first row of hyperintense voxels inferior to VXinf was considered hypothalamic gray matter, and segmented as the ventral border of the hypothalamus (**Figure 1, Panel E)**.

The dorsal border of the hypothalamus in the anterior hypothalamic area (defined as the first slice with ipsilateral interventricular foramen, as in Schindler (Stephanie Schindler et al., 2013) and Bocchetta (Bocchetta et al., 2015)), was aimed to include gray matter in the periventricular region (i.e., to capture the PVN). In the coronal slice where the fornix clearly splits into body and crus, the dorsal border medial to the crus of fornix was considered the dorsal extent of the crus of fornix. The dorsal border lateral to the fornix was defined as in Bocchetta (Bocchetta et al., 2015) (refer to **Table 1**, and **Figure 2**).

**Figure 2.**
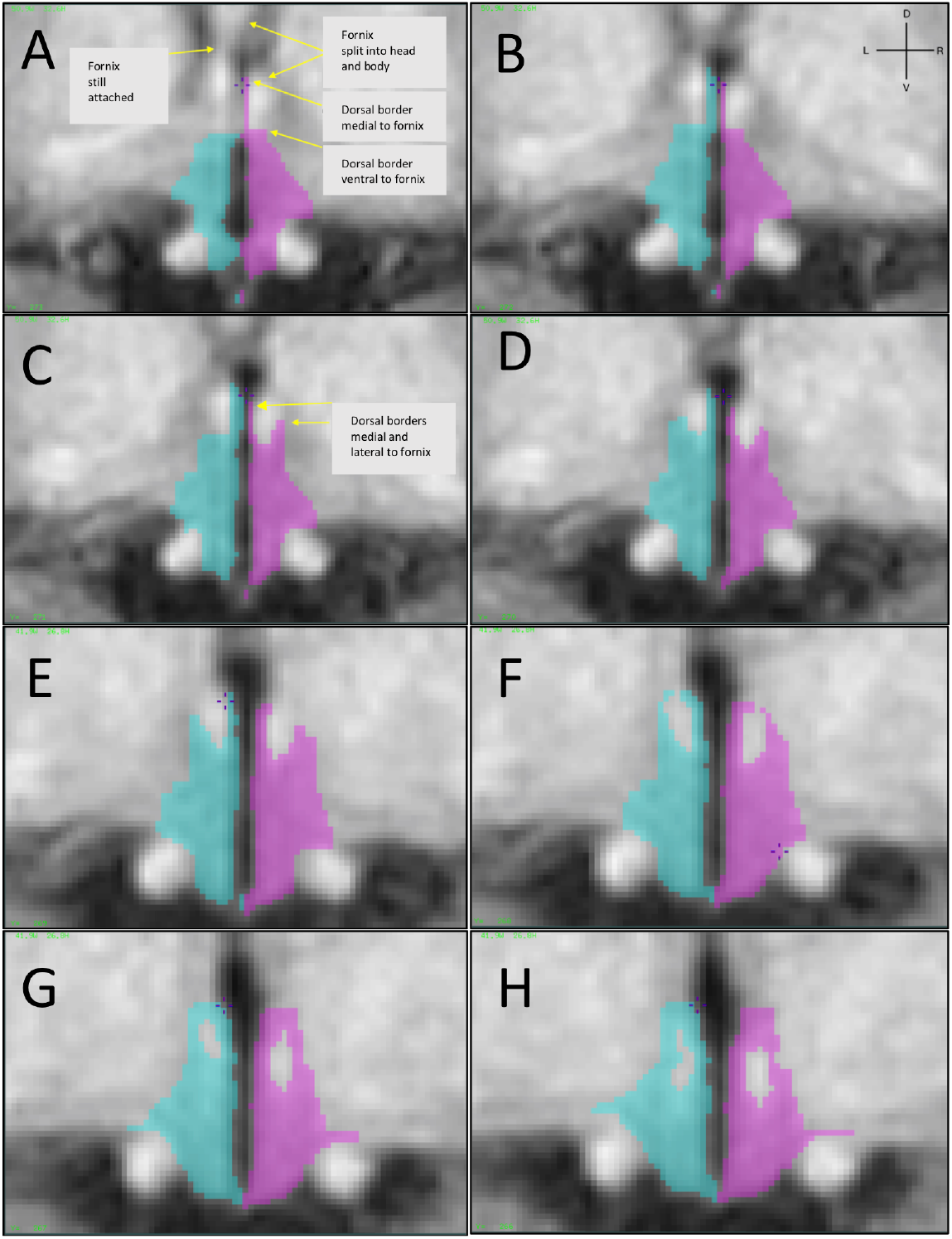
Modifications to Bocchetta when the fornix splits. The dorsal border medial to fornix when the body and crus of fornix split is the superior extent of the fornix (A, purple) and remains the dorsal border up to the hypothalamic sulcus. The lateral border is below the fornix for the first mm (A-B, purple; B-C blue), midway up the fornix for the 2^nd^ mm (C-D, purple; D-E, blue) and above the fornix for the 3^rd^ mm (E-F, purple; F-G, blue). Each voxel = 0.5mm. Orientation is shown in upper right corner of panel B. D: dorsal; L: left; R: right; V: ventral.

The tuberal region was defined from the slice containing the anterior pole of the ipsilateral anteroventral/medial thalamus, to the coronal slice just rostral to the one where the mammillary body was clearly visible (as in Bocchetta et al (2015)). In an effort to capture additional lateral hypothalamic area, the lateral border inferior to the internal globus pallidus was defined as the lateral extent of the optic tract. Because of intermingled gray and white matter in this region, at times voxels above the optic tract would be segmented in a discontinuous manner. In such cases, it was permitted to remove the threshold and fill the gaps by creating a single line of voxels to harmonize the segmentation (**Figure 3**). When the ipsilateral optic tract was more than 10 voxels away from the 3^rd^ ventricular wall in coronal view (or with the appearance of the cerebral peduncle), the lateral extent of the hypothalamus was defined as the medial extent of the optic tract, or the cerebral peduncle which can be easily identified in transverse view.

**Figure 3.**
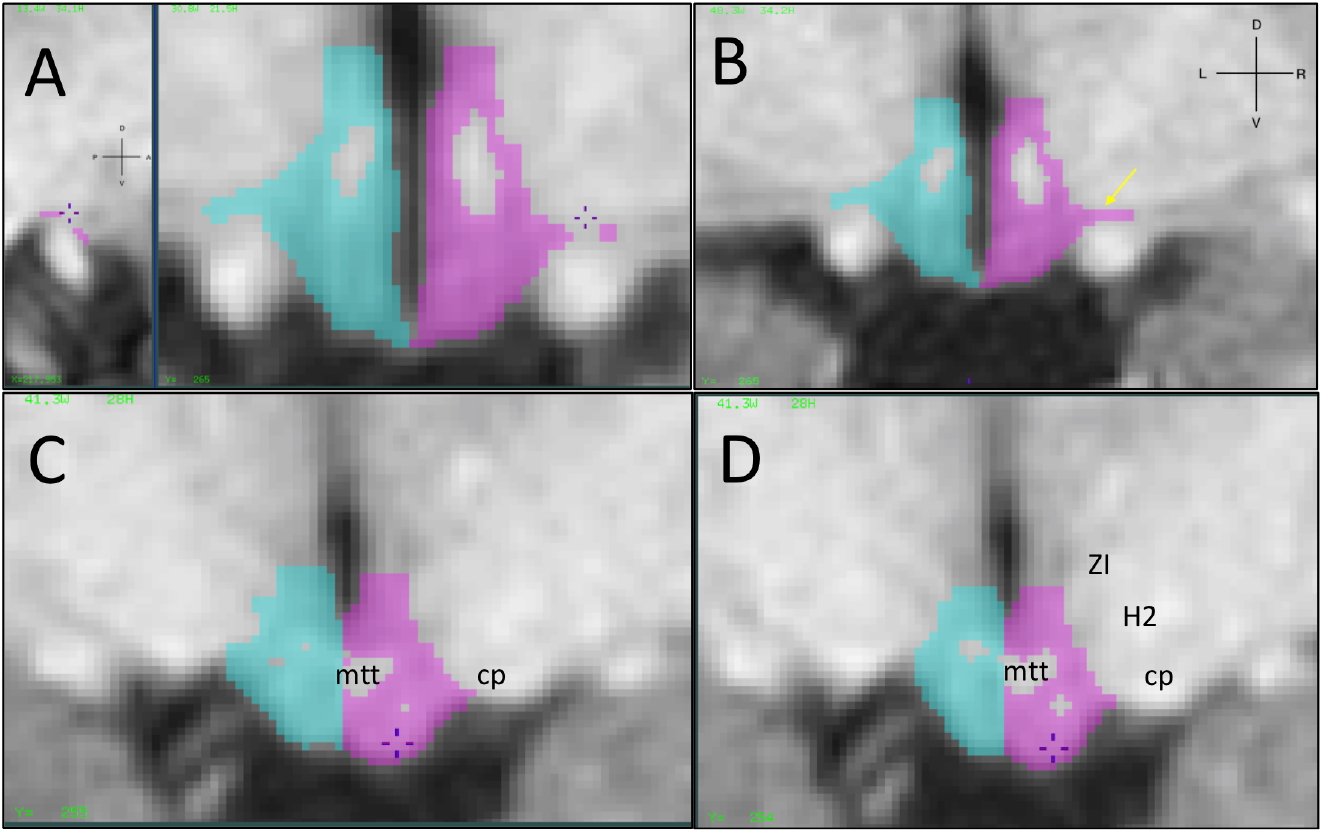
Example of discontinuous segmentation in lateral hypothalamic region with permitted corrections. (A-B), and segmentation in mamillary region (C-D). Lateral border of the lateral hypothalamus can extend to lateral extent of optic tract (A), but threshold may prevent continuous segmentation (unsegmented voxels below crosshair in A). Inclusion of the voxels is permitted by removing the threshold (B). C-D) Note that in the mamillary region, the threshold allows exclusion of the mammillothalamic tract (mtt). Lateral borders are the zona incerta (Zl), fields of forel (H2) and cerebral peduncle. The dorsal border in panel D is an arbitrary horizontal line drawn from superior-medial extent of Zl. Orientation is shown in upper right corner of panel B. D: dorsal; L: left; R: right; V: ventral. Each voxel = 0.5mm.

The mammillary region included the sections that spanned the rostral to caudal extent of the ipsilateral mammillary body. The protocol follows the rules of Bocchetta (Bocchetta et al., 2015) in that regard by using the hypothalamic sulcus as the dorsal border, except a more liberal dorsal border was applied when the hypothalamic sulcus was no longer clearly visible, by drawing an arbitrary line from the ventral medial portion of the zona incerta/field of H2. The mamillothalamic tract was not used as a border, but was excluded from the segmentation by applying the previously established threshold (**Figure 3**).

The hemispheric split is determined by navigating to a section just superior to the mamillary bodies in transverse view, and drawing a straight line that divided the 3rd ventricle and cerebral aqueduct.

##### Pituitary stalk

The dorsal border of the pituitary stalk was defined as the ventral border of the hypothalamus at the level of the stalk. The pituitary stalk is located on the mid-sagittal section of the hypothalamus, and sagittal view was used to follow the course of the stalk and to exclude the optic tract. Segmentation occurred from the superior to inferior extent in the transverse view using the same gray matter contrast intensities that were set for the hypothalamus (see *Setting the Image Contrast* above). Segmentation continued until the stalk was no longer clearly distinguished from surrounding pituitary tissue, which is considered the junction of the pituitary stalk and pituitary gland, similar to the approach defined in Satogami et al. (Satogami et al., 2010).

We verified the segmentation of the hypothalamus in all three orientations, and removed the threshold where necessary, and in this way completed the segmentation surrounding the fornix, and of the lateral hypothalamus (**Figure 3**).

#### Hypothalamus parcellation

For the hypothalamic subregion parcellation, we defined preoptic, periventricular and lateral hypothalamic regions. For completeness, we specified parcellation rules for segmentation of the mammillary bodies, and then classified the remaining unparcelled segments into the anterior, tuberal, and posterior hypothalamic regions. At this point, no additional segmentation was permitted, we restricted ourselves to an allocation of existing labels to hypothalamic subregions. In all cases, we applied the rule that made the most anatomical sense. A summary of parcellation boundaries is provided in Table S1.

##### Preoptic region

The preoptic area (POA) included all sections that were included as anterior preoptic area in the sections anterior to the continuous anterior commissure (as described above), with a maximum lateral border defined as a straight line drawn from the ventral medial edge of the septum or fornix, to the voxel just medial to the “bulge” of the optic tract (VX0; **Table S2 and Figure 4 panel A**). For sections with continuous anterior commissure superior to the 3^rd^ ventricle ipsilaterally, the lateral boundary of the preoptic region was defined as a straight line drawn from a voxel just medial to the appearance of the “bulge” of the optic tract (VX1, **Table S2 and Figure 4 panel B**) to the lateral most extent of the fornix. The ventral border was defined as an arbitrary straight line drawn from the cerebral exterior (**Figure 4 panel C)**. The last section of the POA contained anterior commissure superior to the 3^rd^ ventricle (**Figure 4 panel D)**.

**Figure 4.**
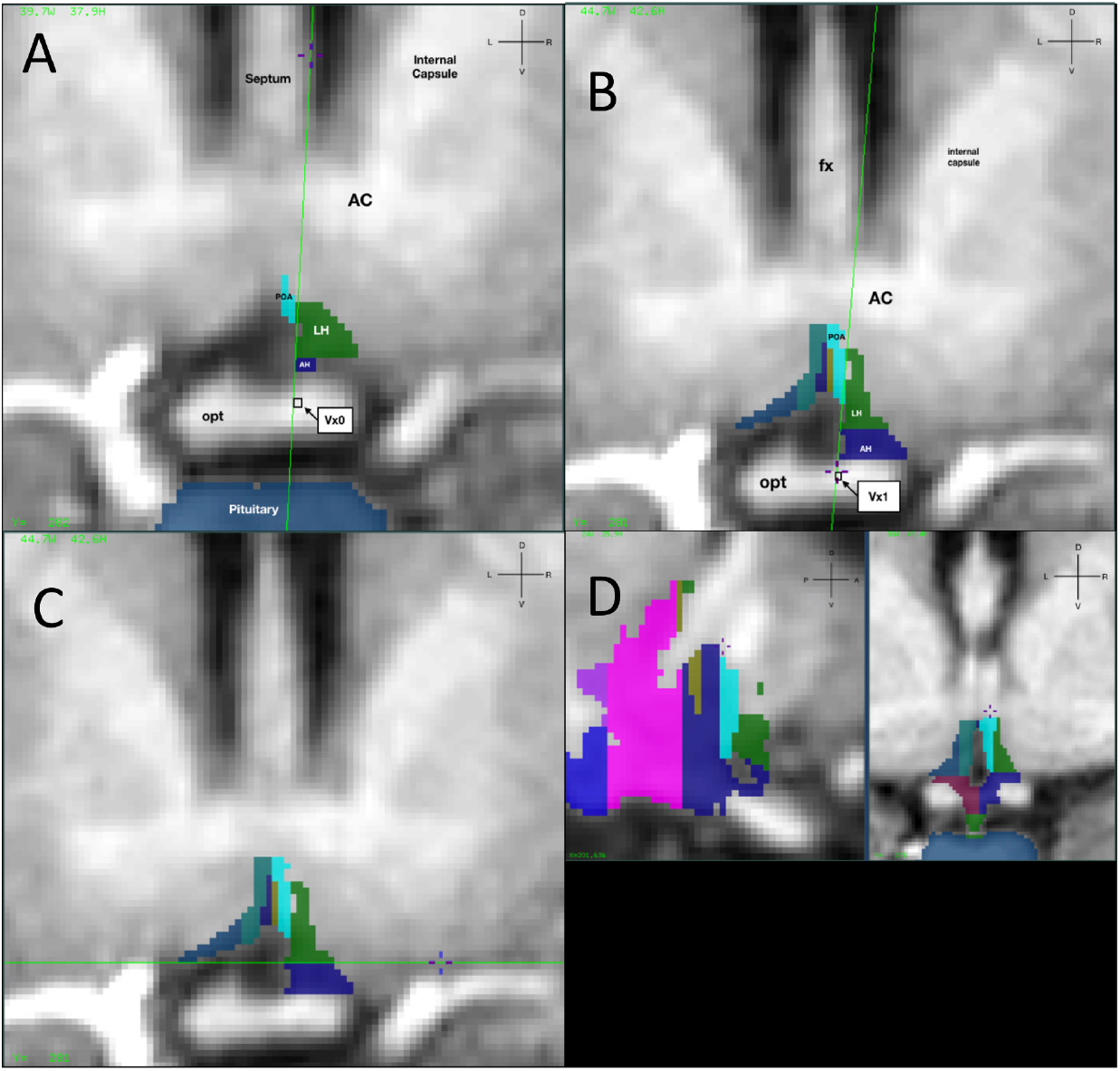
Parcellation of the preoptic region. Refer to right side segmentation for all panels. A. The voxel just medial to the bulge of the optic tract (VxO, shown in the blue box) is determined on sections anterior to the continuous anterior commissure. The lateral border of the preoptic region (POA) is defined as a straight line drawn from the medial edge of the septum or fornix (crosshair), to the voxel just medial to the “bulge” of the optic tract. B. For sections with continuous anterior commissure superior to the 3^rd^ ventricle ipsilaterally, the lateral boundary of the preoptic region is defined as a straight line drawn from a voxel just medial to the appearance of the “bulge” of the optic tract (VXl) to the lateral most extent of the fornix. C. The ventral border is an arbitrary straight line drawn from the cerebral exterior. D. The last section of the POA is that which contains anterior commissure superior to the 3^rd^ ventricle, which can be verified using sagittal plane (see voxel under cross hair in sagittal (left) and coronal (right) planes). AH: Anterior hypothalamus. LH: Lateral hypothalamus. Opt: Optic tract. Orientation is shown in upper right corner of panels. D: dorsal; L: left; R: right; V: ventral. Each voxel = 0.5mm.

##### Periventricular region

Parcellation of the periventricular region began on the coronal section with continuous anterior commissure ipsilaterally, was always bordered by the 3^rd^ ventricle medially, and the ventral border was defined as an arbitrary horizontal line drawn from the lateral cerebral exterior. Navigating from anterior to posterior in the coronal plane, the dorsal and lateral borders were first determined by selecting a voxel that we called VX3v (**Table S2**), defined as the voxel at the midpoint of the dorsal 3^rd^ ventricle, or the voxel at the dorsal hypothalamus that separated the two hemispheres, selecting the most superior of the two, and drawing a straight line to VX1 (see *preoptic region*) (**Figure 5 panel A)**. Note that if the periventricular parcellation overlapped entirely with that of the POA parcellation in rostral sections (i.e., just before the disappearance of the anterior commissure), we assigned the parcellation as periventricular.

**Figure 5.**
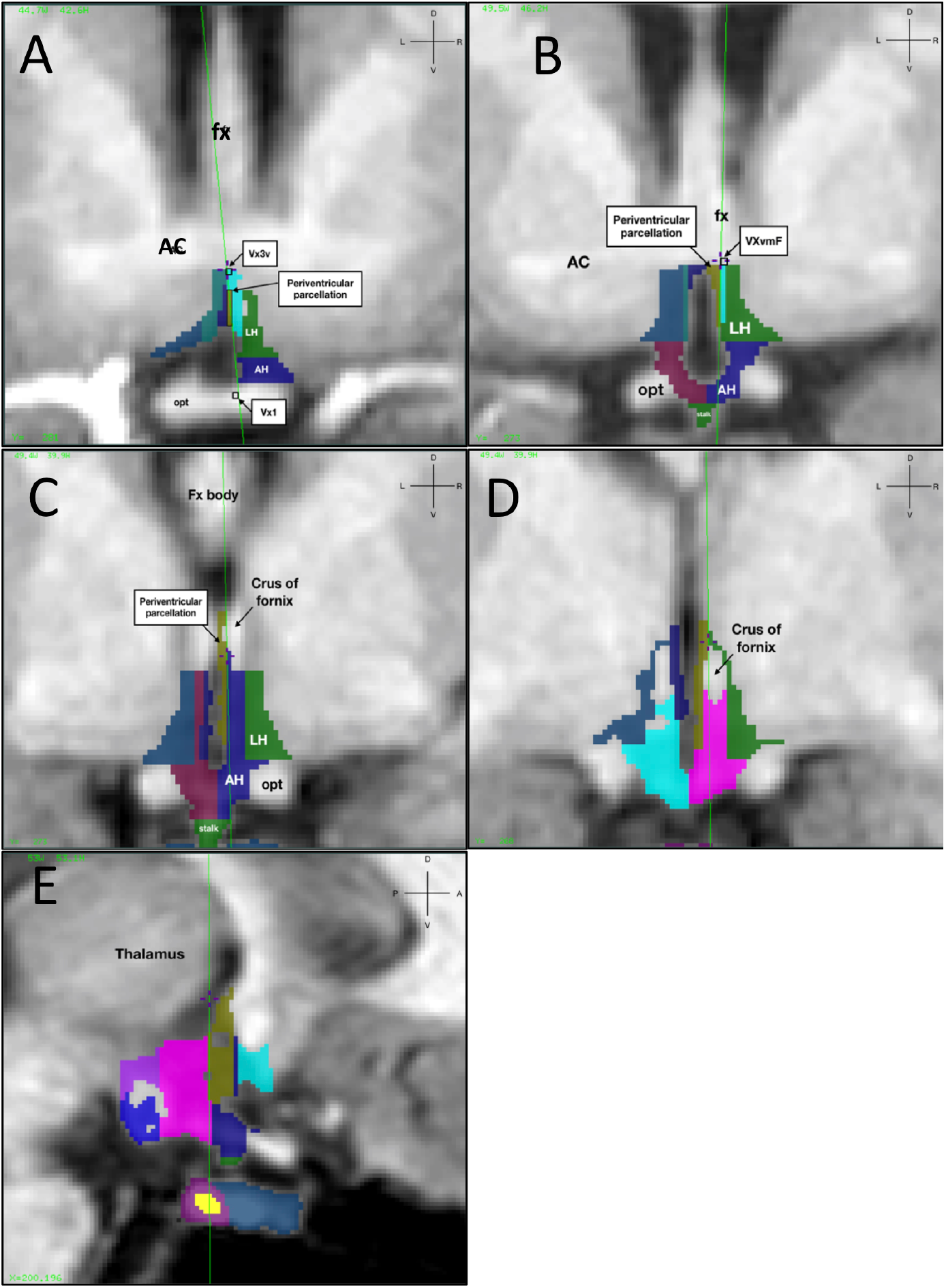
Parcellation of the periventricular region. Refer to right hemisphere for all panels. **A.** Navigating from anterior to posterior, the dorsal and lateral borders in the preoptic region are determined by a straight line drawn from a voxel at the midpoint of the dorsal 3^rd^ ventricle or the dorsal hypothalamus, selecting the point that separates the two hemispheres (VX3v, as shown at crosshair) to VXl (refer to definition in preoptic region). **B.** Once the AC disappears and the fornix emerges just superior to the 3^rd^ ventricle and before the fornix splits into body and crus, the dorsal and lateral borders of the periventricular region are determined by an arbitrary line that connects a voxel at the ventromedial extent of the ipsilateral fornix (VXvmF; crosshair) to VXl (refer to definition in preoptic region) or the voxel at the lateral extent of the infundibular stalk. **C.** On the section where the fornix is split into body and crus on the coronal section or once the infundibular stalk is no longer visible, the dorsal border is the dorsal extent of the fornix, and the lateral border is defined by a straight vertical line drawn from the segmented voxel at the most superior point of the medial extent of the fornix. **D.** When the crus of fornix is no longer touching the 3^rd^ ventricle as it appears to move laterally, the lateral border dorsal to the fornix is the most lateral voxel segmented on the medial extent of the fornix, whereas the lateral border ventral to the fornix is the medial extent of the fornix. **E.** The last section for segmenting the periventricular region is the one just before the thalamus is clearly visible (clicking at the lateral extent of the periventricular parcel), cross-checking in sagittal. Periventricular parcellation is shown in brown. AC: Anterior commissure; AH: Anterior hypothalamic parcel; LH: lateral hypothalamic parcel; opt: optic tract. Fx: fornix. Orientation is shown in upper right corner of panels. D: dorsal; L: left; R: right; V: ventral. **Each voxel =** 0.5mm.

Once the AC disappeared and the fornix emerged just superior to the 3^rd^ ventricle and before the fornix split into body and crus (refer to plates 34-35 in (Mai et al., 2015), the dorsal and lateral borders of the periventricular region were determined by an arbitrary line that connected a voxel at the ventromedial extent of the ipsilateral fornix (VXvmF, **Table S2)** as shown in **Figure 5 (Panel B**) to VX1 or the voxel at the lateral extent of the infundibular stalk, or if neither were visible, a vertical line. On the section where the fornix is split into body and crus on the coronal section or once the infundibular stalk was no longer visible (refer to plate 26 in (Mai et al., 2015)), the dorsal border was the dorsal extent of the crus of fornix, and the lateral border was defined by a straight vertical line drawn from the segmented voxel at the most superior point of the medial extent of the fornix (**Figure 5 Panel C)**. Continuing in the posterior direction, when the crus of fornix was no longer touching the 3^rd^ ventricle as it appeared to move away laterally (refer to Plate 37 in (Mai et al., 2015)), and which we objectively defined as two or more columns of voxels segmented medial to the fornix in its superior-inferior extent, the dorsal border of the periventricular region was defined as the dorsal extent of the fornix or the hypothalamic sulcus (as determined by initial segmentation), and the lateral border dorsal to the fornix was defined as the most lateral voxel segmented on the medial extent of the fornix, whereas the lateral border ventral to the fornix was defined as the medial extent of the fornix (**Figure 5 Panel D**). The last section for segmenting the periventricular region was defined as the one just before the thalamus became clearly visible (with cross-checking in sagittal orientation) (**Figure 5 Panel E**). Note that the parcellation of one extra column of voxels was permitted if it allowed a continuous parcellation with the previous PVN section.

##### Lateral hypothalamus

Parcellation of the lateral hypothalamic region began on the first section containing anterior preoptic area (**Table 1**). The medial border was defined as the preoptic parcellation, and included all hypothalamic segmentation lateral to the preoptic parcel. The ventral border was defined as an arbitrary horizontal line drawn from the lateral cerebral exterior (**Figure 6 Panel A)**.

**Figure 6.**
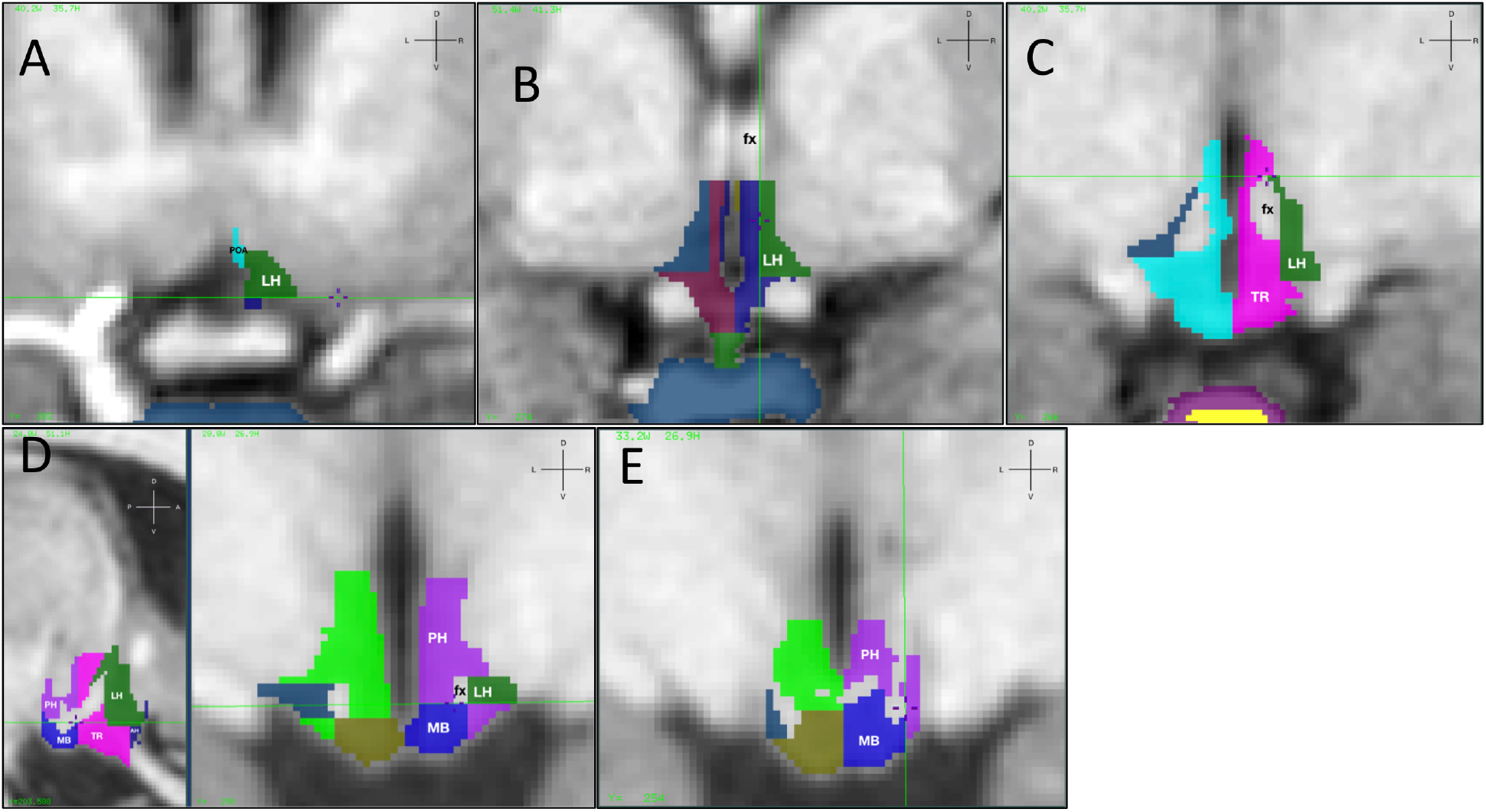
Parcellation of the lateral hypothalamus (LH). Refer to hypothalamus segmentation on right. **A.** Parcellation begins on the first section containing anterior preoptic area. The medial border is the preoptic parcellation, and includes all hypothalamic segmentation lateral to the preoptic parcel. The ventral border is an arbitrary horizontal line drawn from the cerebral exterior. **B.** Posterior to the anterior commissure and with the emergence of the fornix just superior to the 3^rd^ ventricle, the medial border is an arbitrary vertical line drawn from the lateral extent of the fornix, or the periventricular region and the parcellation includes all segmented hypothalamus lateral to the medial border. **C.** When the crus of fornix is no longer touching the 3^rd^ ventricle, up to the section just prior to the emergence of the thalamus, the dorsal and medial borders include all segmentation lateral to the periventricular region, or, as shown, an arbitrary horizontal line drawn at the superior extent of the fornix. **D.** Once the mammillary bodies appear, the medial border is the fornix, the ventral border is an arbitrary horizontal line drawn from the ventral extent of the fornix, or the most lateral voxel segmented superior to the optic tract, whichever is most inferior, or the cerebral peduncle. The dorsal border is an arbitrary horizontal line drawn from the superior extent of the fornix. The sagittal section is shown on the left side of the panel and the coronal on the right. E. Segmentation of the mamillary bodies using mammillothalamic tract as dorsal boundary. PH: Posterior hypothalamic parcel; LH: Lateral hypothalamic parcel; MB: Mamillary Body; fx: fornix. Orientation is shown in upper right corner of panels. D: dorsal; L: left; R: right; V: ventral. Each voxel = 0.5mm.

Once the anterior commissure disappeared and the fornix emerged just superior to the 3^rd^ ventricle (refer to plates 34-35 in (Mai et al., 2015)), the medial border was defined as an arbitrary vertical line drawn from the lateral extent of the fornix, or the periventricular region and the parcellation included all segmented hypothalamus lateral to the medial border (**Figure 6 Panel B)**. The dorsal border was defined as an arbitrary horizontal line drawn from the superior edge of VXvmF. The ventral border was defined as a horizontal line drawn from the cerebral exterior, or the optic tract. Continuing in the posterior direction, when the crus of fornix was no longer touching the 3^rd^ ventricle as it appeared to move laterally (refer to Plate 37 in (Mai et al., 2015)), up to the section just prior to the emergence of the thalamus, the dorsal border included all segmentation lateral to the periventricular region, or an arbitrary horizontal line drawn from the superior extent of the fornix (**Figure 6 Panel C)**. The medial border was defined as the medial extent on the lateral edge of the ipsilateral fornix, and the ventral border as the most lateral voxel segmented superior to the optic tract.

Once the thalamus appeared (refer to plates 38-39 (Mai et al., 2015)), the medial border followed the fornix or the mammillothalamic tract (selecting the most medial of the two), the dorsal border was defined as a horizontal line drawn from the superior extent of the fornix, or the most lateral voxel segmented superior to the optic tract, selecting the most superior of the two. The ventral border extended from the inferior extent of the fornix to the most lateral voxel segmented superior to the optic tract, selecting the most inferior of the two.

Once the mammillary bodies appeared, the medial border was defined as the fornix, and the ventral border as an arbitrary horizontal line drawn from the ventral extent of the fornix, or the most lateral voxel segmented superior to the optic tract, whichever was most inferior, or the cerebral peduncle. The dorsal border was defined by an arbitrary horizontal line drawn from the superior extent of the fornix (**Figure 6 Panel D)**, and the lateral border was the cerebral peduncle. Segmentation of the lateral hypothalamus ended on the last section containing segmented fornix, or the section just before the mamillary bodies touched the cerebral peduncle laterally.

The mammillary bodies were parcelled using the fornix or mammillothalamic tract (verifying in sagittal view) as the dorsal boundary (**Figure 6 Panel E)**, the lateral border in hierarchical order was defined as the lateral hypothalamus, the lateral extent of the fornix, or the cerebral peduncle. The medial border was defined as the 3^rd^ ventricle or midline, and the ventral border as the cerebral exterior. We used transverse view to confirm lateral boundaries.

##### Anterior hypothalamus, Tuberal region, and Posterior Hypothalamus Parcellation

For completeness, the anterior hypothalamus included all hypothalamic segmentations that were not segmented as preoptic, periventricular or lateral hypothalamus from the first section segmented to the caudal most tip of the segmented infundibulum when attached to the hypothalamus in coronal view, or the caudal extent of the preoptic region for brains where the stalk swung forward.

The tuberal region parcellation consisted of all hypothalamic segmentation between the section just caudal to the infundibulum to the slice just rostral to the one that clearly contained mammillary bodies, and that was not segmented as lateral hypothalamic area or periventricular area.

The posterior hypothalamus parcellation consisted of the hypothalamic segmentation from the rostral to caudal extent of the ipsilateral mammillary bodies that was not segmented as mammillary body, or lateral hypothalamic area.

##### Pituitary Gland

The pituitary gland is easily identifiable on 3T T1-weighted images, in the mid-sagittal plane, just inferior to the hypothalamus. The pituitary gland is bordered by the sphenoid sinus anteriorly and ventrally, the cavernous sinuses laterally, the dorsum sellae posteriorly, and the diaphragm sellae dorsally (Klomp et al., 2012). Typically, a characteristic feature of the pituitary is a posterior bright spot, which is thought to represent vasopressin cells (Kitagawa et al., 2017; Lee et al., 2001; Sato et al., 1995).

Because the hyper-intensity of the pituitary gland differs from that of the hypothalamus, we adjusted the contrast setting prior to segmenting the pituitary (see ‘*Setting the Image Contrast: Pituitary’).* Prior to beginning the segmentation, we adjusted the threshold (lower limit remained the same as the hypothalamus, and the upper limit value entered at the start of segmentation was the image contrast maximum value).

Segmentation of the pituitary gland began in the sagittal view. On 0.5mm isotropic images, an apparent drop in signal intensity can be observed within the pituitary gland on the T1 weighted image, with a corresponding hyperintense signal on T2. Our identification of a border between anterior and posterior pituitary gland on T2 images was inspired by that observed in animals (Theunissen et al., 2010) using high-field MRI. We defined the hypointense signal on T1 and hyperintense signal on the T2, as a “line” that separated the anterior and posterior pituitary glands (**Figure 7, Panels A and B**). Because the *pars intermedia* is a third lobe positioned between the anterior and pituitary gland, and is sometimes considered to belong to the anterior pituitary gland in humans (Satogami et al., 2010), we included only one line of voxels of that hypo-intense signal as the posterior pituitary gland. This line was identified at the beginning of the segmentation.

**Figure 7.**
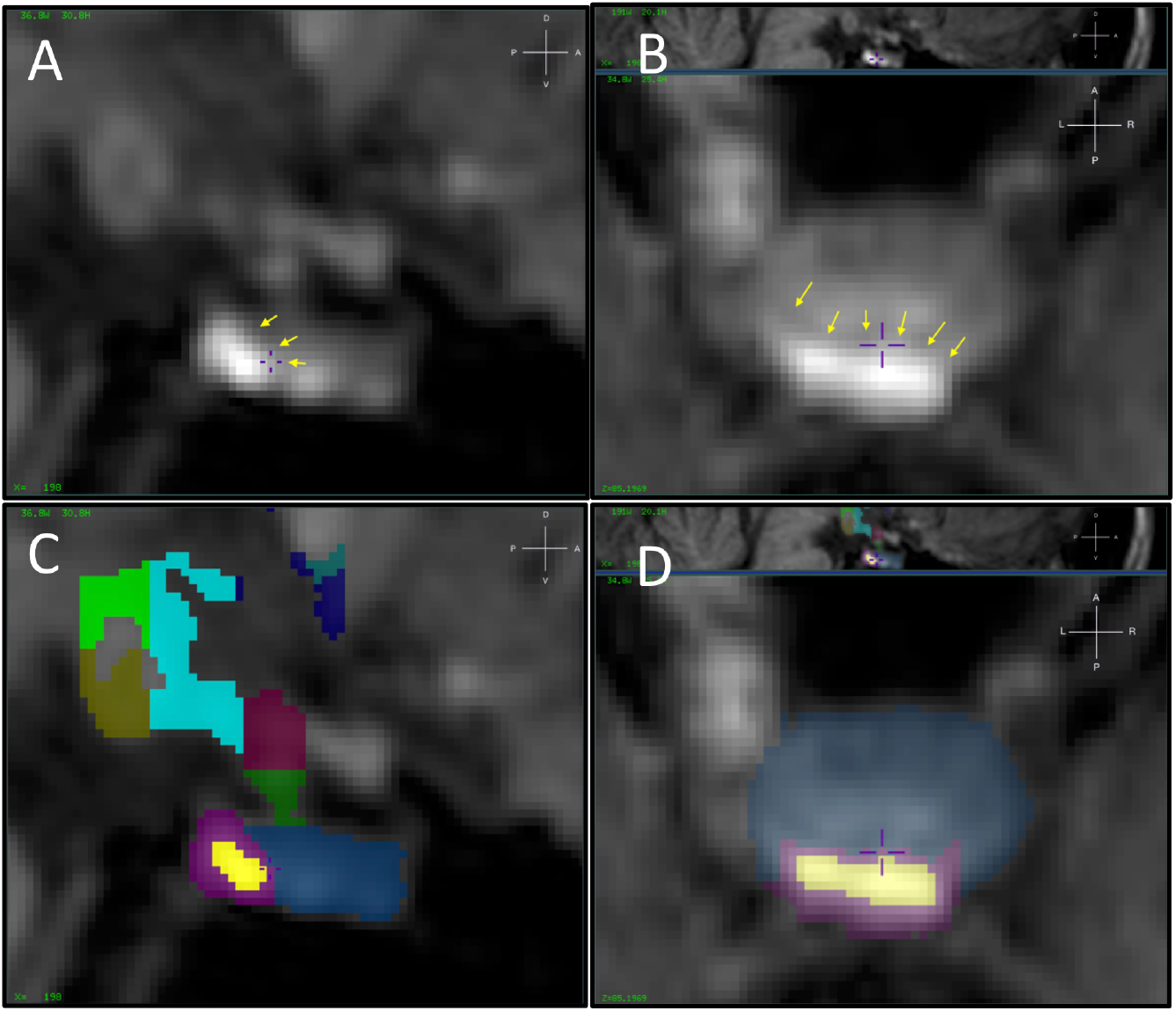
Segmentation of the pituitary gland. **A.** The hypointense signal in the posterior region of the gland is identified on the T1 weighted image in sagittal view and is segmented on all sagittal sections where it is clearly identified. **B.** Following its initial segmentation in the sagittal plane, segmentation of the posterior line ten continues in the transverse plane, where it is also identified as a line of hypointense signal in the posterior region. The anterior pituitary segmentation is completed in the coronal plane (not shown). Complete segmentations are shown in the sagittal (panel **C**) and transverse (panel **D**) planes. Orientation is shown in upper right corner of panels. D: dorsal; L: left; R: right; V: ventral. Each voxel = 0.5mm.

First, using the label color of the posterior pituitary gland, we drew a line that divided the anterior and posterior glands on all sagittal images where a separation was clearly visible (**Figure 7 panel A)**. Next, from the most superior extent of the gland and moving in the inferior direction, we segmented the posterior pituitary gland in the transverse plane (**Figure 7 panel B**), on all sections where the hypointense signal was clearly seen. At this point, we filled the label of the posterior pituitary gland.

Next, in the sagittal orientation, we navigated to the midline of the pituitary gland. Using the anterior pituitary gland label, we drew the rostral extent of the anterior pituitary in sagittal view and drew the outline in transverse view. We then completed the segmentation using the coronal orientation, beginning on the most rostral section previously labelled in sagittal view.

Finally, to label the posterior bright spot, we entered the threshold limits and segmented all voxels in the posterior pituitary that exceeded the 75% threshold, as posterior bright spot.

Lastly, we verified the segmentation of the pituitary in all three orientations, harmonizing borders and overall shape of the structure. In cases where single voxels were not segmented but were clearly part of the structure, we removed the threshold to be able to include those. **Figure 7** shows the complete segmentation in the sagittal (panel C) and transverse (panel D) planes.

#### Reliability and Validity Assessment

A subset of scans were randomly selected from the pool of scans, for reliability analyses.

Intrarater reliability was assessed by a single rater (SLJ) by segmenting 5 subjects two times, with at least one-month interval between segmentations. Interrater reliability was assessed by three different raters (SLJ, CA, MD) segmenting 3 subject scans two times with at least one month interval between segmentations. All raters were blind to sex, and two of the three raters (CA, MD) to the expected sex differences.

For validation purposes, sex differences in mean volume were expected in whole hypothalamus (Makris et al., 2013) as well as whole pituitary gland (MacMaster et al., 2007; Takano et al., 1999; Wong et al., 2014). As a validation of the anterior and posterior divisions, we expected that the posterior pituitary gland occupied in relation to the total gland would be approximately 20-30%, as reported histologically (Hong et al., 2016).

### Statistical Analyses

Dice kappa coefficients were calculated for voxel overlap (Feuerman & Miller, 2008) to test inter- and intra-rater reliability.

All variables were verified for normality. Testosterone values were log transformed because they were not normally distributed. Although estradiol levels were positively skewed, the log transformation did not improve the normality of the distribution; we thus decided to use raw estradiol levels. All other variables were normally distributed.

Independent t-tests were used to assess steroid hormone levels, as well as the volumetric sex differences hypothesized for hypothalamus and pituitary. Paired-samples t-tests were used to test for hemispheric differences. ANCOVAs were used to assess sex differences in hypothalamic and pituitary volume, covarying for those hormones where a sex difference was detected, or where the hormone was associated with hypothalamic or pituitary measures. Cohen’s *d* and partial eta squared (ηp^2^) are reported as measures of effect size.

Finally, because the hypothalamus and pituitary gland are involved in the production of sex steroid hormone levels, the hypothalamus is dense in estrogen (Kruijver et al., 2002, 2003) and androgen (Fernández-Guasti et al., 2000) receptors, and because sex steroid hormone levels have been shown to be associated with brain volume (Herting et al., 2014; Koolschijn et al., 2014), exploratory correlational analyses were conducted to assess associations between hypothalamic and pituitary volumes, as well as the associations between these neuroendocrine structures and salivary estradiol and testosterone levels.

## Results

### Descriptive statistics

The mean volume of the left, right, and whole hypothalamus, the pituitary stalk, as well as the anterior and posterior lobes of the pituitary gland, the total pituitary gland, the pituitary bright spot, and raw steroid hormone levels are shown in **Table 2**. Total brain volume, with and without CSF, was larger in men than in women. Men had significantly higher salivary testosterone levels than women.

**Table 2.** Descriptive Statistics and Sex Differences in Hypothalamus, Pituitary, Total Intracranial and Brain Volumes, and Salivary Hormone Levels.

| Structure | Total Sample (n=31) |  | Men (n=17) |  | Women (n=14) |  | p-value<br>Sex Difference | Cohen's d for the<br>sex difference |
| --- | --- | --- | --- | --- | --- | --- | --- | --- |
|  | Mean | SD | Mean | SD | Mean | SD |  |  |
| Hypothalamus left (mm<sup>3</sup>) | 796.98 | 138.57 | 851.00 | 120.64 | 731.39 | 133.98 | **0.014** | 0.938 |
| Hypothalamus right (mm<sup>3</sup>) | 798.40 | 148.52 | 862.29 | 144.32 | 720.82 | 115.97 | **0.006** | 1.081 |
| Hypothalamus Whole (mm<sup>3</sup>) | 1595.38 | 282.85 | 1713.29 | 259.00 | 1452.21 | 248.62 | **0.008** | 1.028 |
| Medial Preoptic Region left (mm<sup>3</sup>) | 25.88 | 12.17 | 28.81 | 13.92 | 22.31 | 8.86 | 0.142 | 0.557 |
| Medial Preoptic Region right (mm<sup>3</sup>) | 24.63 | 12.34 | 28.06 | 14.64 | 20.46 | 7.31 | 0.09# | 0.657 |
| Paraventricular region left (mm<sup>3</sup>) | 46.49 | 28.82 | 51.17 | 36.65 | 40.81 | 14.16 | 0.328 | 0.373 |
| Paraventricular region right (mm<sup>3</sup>) | 39.34 | 24.91 | 44.08 | 32.29 | 33.58 | 9.11 | 0.249 | 0.443 |
| Lateral hypothalamus left (mm<sup>3</sup>) | 184.59 | 50.16 | 193.91 | 55.28 | 173.27 | 42.35 | 0.261 | 0.419 |
| Lateral hypothalamus right (mm<sup>3</sup>) | 170.20 | 49.61 | 179.68 | 57.61 | 158.68 | 36.51 | 0.247 | 0.436 |
| Anterior hypothalamic area left (mm<sup>3</sup>) | 87.50 | 47.22 | 98.54 | 53.25 | 74.08 | 36.08 | 0.154 | 0.538 |
| Anterior hypothalamic area right (mm<sup>3</sup>) | 92.05 | 51.93 | 108.01 | 54.48 | 72.67 | 42.81 | 0.058# | 0.721 |
| Tuberal hypothalamic area left (mm<sup>3</sup>) | 251.04 | 59.32 | 257.00 | 59.72 | 243.80 | 60.23 | 0.547 | 0.220 |
| Tuberal hypothalamic area right (mm<sup>3</sup>) | 265.08 | 62.19 | 273.35 | 62.12 | 255.04 | 63.08 | 0.424 | 0.293 |
| Posterior hypothalamic area left (mm<sup>3</sup>) | 122.20 | 43.05 | 136.21 | 39.20 | 105.19 | 42.63 | **0.044** | 0.758 |
| Posterior hypothalamic area right (mm<sup>3</sup>) | 116.00 | 49.00 | 129.18 | 51.52 | 100.00 | 42.07 | 0.100 | 0.620 |
| Mammillary body left (mm<sup>3</sup>) | 77.32 | 19.49 | 81.94 | 18.12 | 71.71 | 20.28 | 0.149 | 0.532 |
| Mammillary body right (mm<sup>3</sup>) | 84.34 | 23.34 | 87.75 | 23.47 | 80.21 | 23.36 | 0.380 | 0.322 |
| Stalk (mm<sup>3</sup>) | 27.67 | 10.09 | 29.96 | 10.21 | 24.90 | 9.58 | 0.169 | 0.511 |
| Pituitary anterior (mm<sup>3</sup>) | 684.36 | 134.98 | 648.40 | 140.79 | 728.03 | 117.91 | 0.103 | -0.613 |
| Pituitary posterior (mm<sup>3</sup>) | 211.11 | 42.52 | 223.24 | 45.87 | 196.38 | 34.01 | 0.080 | 0.665 |
| Pituitary bright spot (mm<sup>3</sup>) | 47.09 | 30.64 | 46.78 | 33.94 | 47.46 | 27.36 | 0.952 | -0.022 |
| Pituitary Whole (mm<sup>3</sup>) | 895.47 | 144.64 | 871.64 | 151.83 | 924.40 | 135.14 | 0.320 | -0.367 |
| Total Brain Volume + CSF (cm<sup>3</sup>) | 1350.96 | 90.80 | 1389.84 | 89.46 | 1303.75 | 69.28 | **0.006** | 1.076 |
| Total Brain Volume - CSF (cm<sup>3</sup>) | 1150.11 | 82.25 | 1182.53 | 78.72 | 1110.74 | 70.30 | **0.013** | 0.962 |
| Testosterone (pg/mL) | 88.68 | 58.93 | 125.66 | 49.38 | 39.38 | 23.65 | **<0.001** | 2.229 |
| Estradiol (pg/mL) | 1.41 | 0.80 | 1.65 | 1.01 | 1.12 | 0.23 | 0.058 | 0.724 |

Note. <sup>a</sup>N=31; except for Testosterone (N=28, 16M 12W); Estradiol (N=29, 16M, 13W).
T-tests for sex difference was performed on the log transformed testosterone levels.
CSF: cerebrospinal fluid.

### Reliability

Dice kappa overlap intra- and inter-rater reliability coefficients are shown in **Table 3**. Both the inter-rater and the intra-rater Dice overlap coefficients were approximately 0.83 for left, right and whole hypothalamus, and approximately 0.90 for the pituitary gland and its subregions, including the small pituitary bright spot (M=47.09mm^3^). The pituitary stalk also had acceptable intra- and inter-rater reliability of 0.79, despite its small volume (M=27.67mm^3^). Mean intra- and inter-rater reliability coefficients for the POA, PVN, and LH parcellations ranged between 0.58-0.67.

**Table 3.**
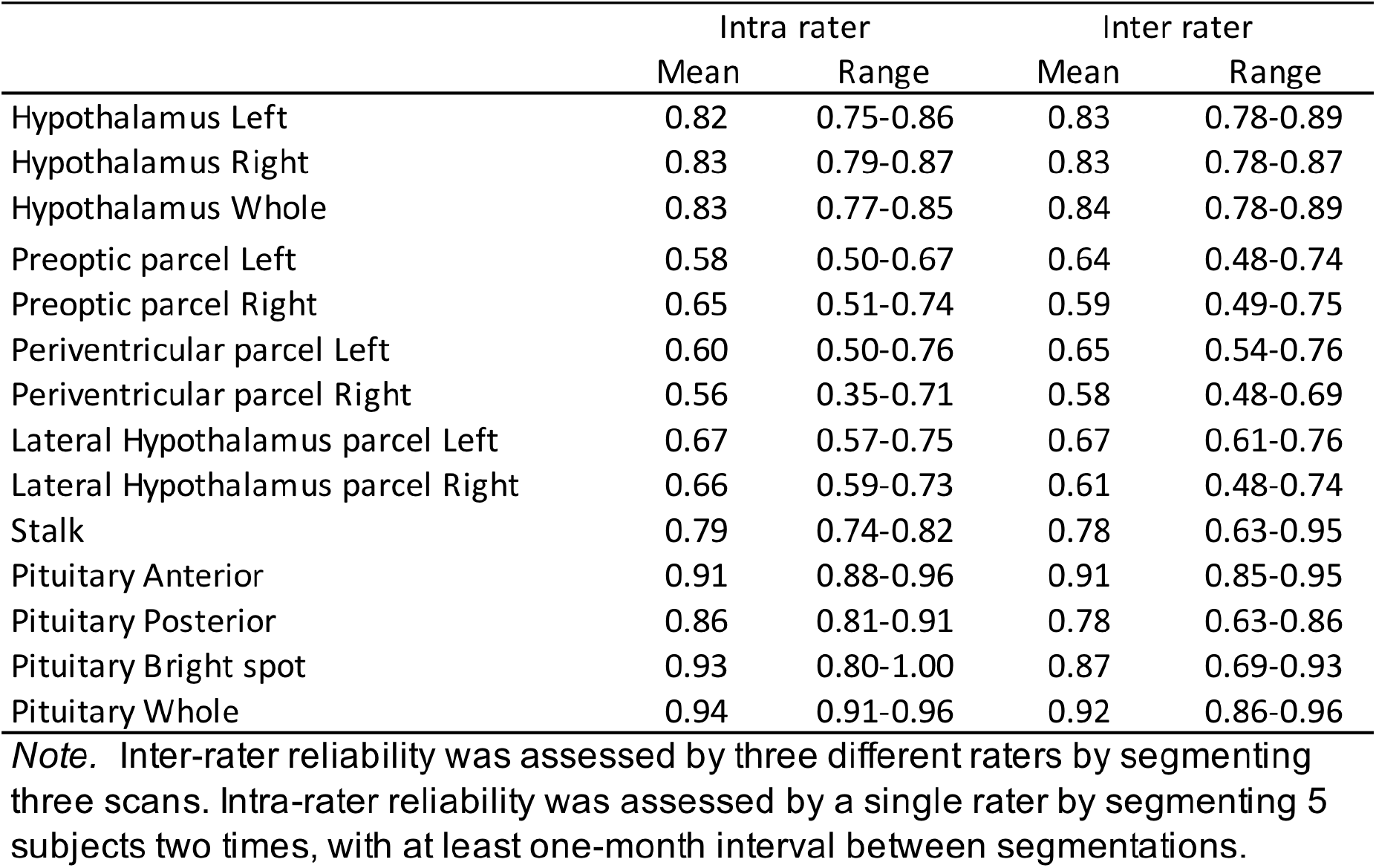
Intra- and Inter-rater Dice Kappa Reliability Coefficients.

### Validity

Independent t-tests were used to assess sex differences on whole hypothalamus, and paired t-tests on each hemisphere. As expected, total hypothalamic volume as well as both hemispheres were significantly larger in males than in females (**Table 2**).

We also tested whether left and right hemisphere volumes differed. No hemispheric differences in hypothalamic volume were detected in the whole sample, *t*(30)=0.157, p=0.876, Cohen’s d=0.03, nor when stratified by sex (women, *t*(13)=1.257, p=0.231, Cohen’s *d*=0.34; men, *t*(16)=0.767, p=0.454, Cohen’s *d*=0.19), consistent with others (Schindler et al., 2013).

Total pituitary volume did not differ between men and women, nor did total anterior pituitary volume, however men tended to have a larger posterior pituitary gland than women (**Table 2**), consistent with Takano et al., (Takano et al., 1999).

The posterior pituitary gland comprised 23.99% [range: 15.39-38.23%] of the total pituitary volume, consistent with histological studies.

Hypothalamic parcellations. The right medial preoptic region was nominally larger in men than in women, however this difference did not attain statistical significance (**Table 2**). No other sex differences were detected.

**Table 4** shows exploratory correlational analyses between hypothalamic and pituitary structures, and sex steroid hormones. For women, salivary estradiol levels were moderately correlated with TBV+CSF and TBV-CSF (r’s=0.57, p<0.05), but not with hypothalamic or pituitary volumes. No associations were detected in men. Salivary testosterone levels were not correlated with TBV+CSF, TBV-CSF, nor with the whole hypothalamus or pituitary gland, or with the subregions of the pituitary.

**Table 4.**
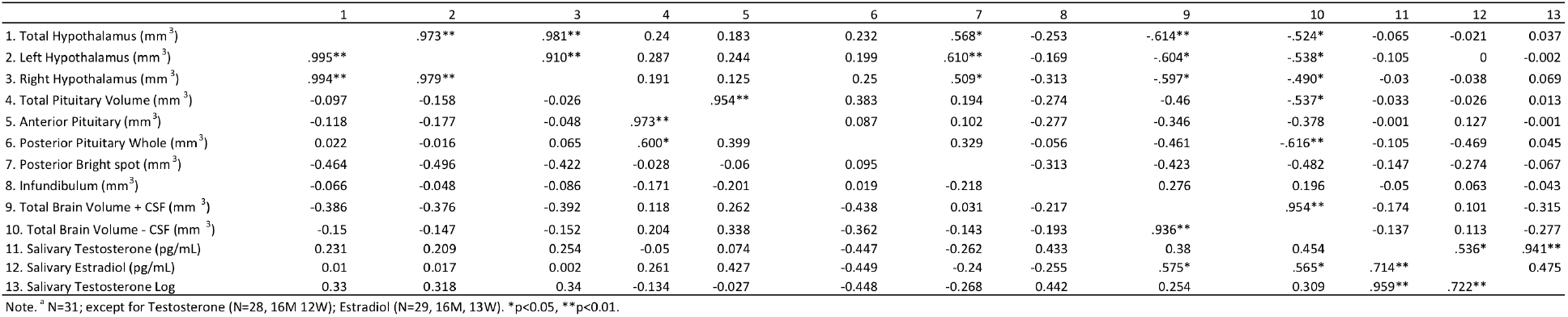
>Pearson Correlation Table Between Hypothalamus, Pituitary, and Salivary Hormones for boys (above diagonal) and girls (below diagonal).

In women, log transformed salivary testosterone levels correlated with total PVN (left+right hemispheres) (r=0.643, p=0.24), which was driven by the left PVN (r=0.663, p=0.019; right PVN, r=0.447, p=0.145). A closer examination of that association suggested that it may have been driven by one woman with higher T levels (with a raw value of 108.7 pg/mL). When she was removed from the analysis, the association was weakened for total PVN (r=0.584, p=0.059), and the association was instead significantly related to right PVN volume (r=0.714, p=0.014), not left PVN (0.414, p=0.205). No other associations between hormone levels and hypothalamic parcellations were detected in females or in males (data not shown).

### ANCOVAs for sex differences controlling for estradiol and testosterone

A sex difference in salivary testosterone levels was detected (**Table 2**). Thus, we tested whether the sex difference in hypothalamic volume was maintained when covarying for testosterone, but this abolished the sex differences. The sex difference was no longer detected in left (p=0.543, ηp^2^ = 0.015), right (p=0.437, ηp^2^=0.024) or total (p=0.480, ηp^2^=0.020) hypothalamic volume, nor in the left posterior hypothalamic area (p=0.222, ηp^2^=0.059).

Similarly, the trend in sex difference was no longer detected in the right medial preoptic area (p=0.475, ηp^2^=0.021), or posterior pituitary volume (p=0.200, ηp^2^=0.065) after controlling for salivary testosterone. No other regions attained significance when controlling for testosterone.

Salivary estradiol levels tended to be higher in men than in women (**Table 2**), and estradiol levels were positively correlated with brain volume for women but not men (**Table 4**). Thus, we tested for sex differences in hypothalamic and pituitary volume, covarying for estradiol. We found that the sex differences (shown in **Table 2**) were maintained in total hypothalamic (p=0.030, ηp^2^=0.169), and right hypothalamic volume (p=0.019, ηp^2^=0.194), and the sex difference was reduced to a trend for left hypothalamic volume (p=0.054, ηp^2^=0.135).

Within the hypothalamic parcels, right MPA was now significantly larger in men than women (p=0.045, ηp^2^=0.146). A trend emerged on the right PVN, which tended to be larger in men than in women (p=0.085, ηp^2^=0.110). The larger left posterior hypothalamic region in men compared to women was reduced to a trend (p=0.051, ηp^2^=0.138). No sex differences were detected on any other hypothalamic parcel. Covarying for estradiol revealed that men had a larger posterior pituitary than women (p=0.022, ηp^2^=0.186), whereas women tended to have a larger anterior pituitary volume (p=0.079, ηp^2^=0.114). Total pituitary volume did not display a sex difference when controlling for estradiol (p=0.319, ηp^2^=0.038).

## Discussion

The manual segmentation protocol for hypothalamus and pituitary segmentation presented here uses uniform and standardized T1 weighted images acquired on a 3T MRI scanner, for comprehensive morphometric study of the hypothalamic-pituitary axes. It was shown to possess good inter- and intra-rater reliability. Moreover, by cross-checking the upsampled T1 weighted images (0.5 mm isotropic) with the high resolution T2 weighted images (acquired at 0.4x0.4 mm in plane), it extends the existing literature by allowing refinement of hypothalamic boundaries, and by defining a parcellation of the hypothalamus in its medial-lateral extent. In addition, the protocol includes the pituitary stalk and a detailed segmentation of the anterior and posterior pituitary gland, which are anatomically and functionally distinct endocrine-regulating regions. Furthermore, adjusting for salivary sex steroid hormone levels affected the magnitude of the sex differences in hypothalamic and pituitary volumes and their substructures. Thus, this protocol provides a novel, unified, comprehensive structural analysis of these neuroendocrine structures.

A feature of this protocol involves an objective definition of the gray matter intensity for segmentation. The hypothalamus and pituitary are bordered by extreme hypo- and hyper- intensity signals. Hypointensity signals arise from the cerebral spinal fluid in the 3^rd^ ventricle comprising the medial, and the subarachnoid space for the ventral borders of the hypothalamus, and the sphenoid space which surrounds the pituitary. Hyperintensity signals arise from large white matter tracts such as the AC, Fx, Opt for the hypothalamus, and the posterior bright spot and carotid arteries for the pituitary gland. Although we determined the contrast in our images to lie at the 40% and 75% range on our standardized, and intensity normalized T1-weighted images acquired using MP-RAGE, we suggest that research groups determine the appropriate range for their respective datasets which may depend on acquisition parameters, scanner model and software, and/or variation in subjects. However, we note that these proportions appear to be at least valid in other datasets acquired in our lab using similar MP-RAGE parameters, of healthy subjects varying in age from 11-16 years of age (Jones, S.L., Anastassiadis, C., Dupuis, M., Devenyi, G. A., McKee, K., Laplante, D.P., King, S., Chakravarty, M.M., Pruessner, J.C., *unpublished observations*). We also note that even in scans with movement artifact, the hypothalamic segmentation rules could be applied consistently, and had good reliability. However, the parcellation tended to be more difficult on scans with movement artifacts, particularly for the periventricular region which lines the 3^rd^ ventricle and is bordered by the hyperintense fornix. As such, we recommend to verify movement artifacts at the time of image acquisition and to allow extra time in the scanner to repeat the respective sequences when necessary.

Importantly, one challenge in hypothalamic segmentation is defining the anterior border of the preoptic region, particularly because it is difficult to discern it from septal tissue. Recently, Butler and colleagues (Butler et al., 2014) developed a novel and reliable segmentation protocol of the septal region on T1-weighted images acquired using an MPRAGE sequence at 1x1x1.3mm resolution. A detailed examination of the Butler protocol suggests that there may be overlap between the anterior hypothalamus as described in the current protocol. This is particularly likely given that Butler et al. (Butler et al., 2014) define the inferior boundary of the septum to be the base of the brain in slices anterior to the continuous anterior commissure (as can be seen in Butler et al., (Butler et al., 2014), Figure 2 panels C and D).

Although this is a reasonable and reliable boundary, it is likely to include anterior preoptic tissue that lies along the inferior-ventral border of the diencephalon in those slices (Saper et al., 2004). Most existing protocols consider the first anterior slice of the hypothalamus to be the first slice with continuous anterior commissure, which is a reasonable definition particularly on standard 1mm isotropic resolution images. One exception is the protocol employed by Makris et al. (Makris et al., 2013), that begins segmentation on the section containing the rostral most tip of the anterior commissure and optic chiasm; similarly, our protocol includes the tissue rostral to the continuous anterior commissure, but defines structural characteristics of the lateral anterior commissure on those sections. Because of the higher resolution we acquired in our scans compared to those described in Bocchetta (Bocchetta et al., 2015) (i.e., our T2- weighted images acquired at 0.4mmx0.4mm in plane resolution), we extended the anterior segmentation to include more of the anterior preoptic region, due to its importance in regulating sexual and maternal behaviors, and reproductive physiology (Sisk & Foster, 2004). Importantly, our proposed protocol attempts to reduce segmentation of the septum by defining the superior extent of the anterior preoptic region on those slices anterior to the continuous anterior commissure, based of inferior borders of the anterior commissure which emerge laterally in the coronal slices. This seemed a reasonable geometric rule given that it continues to rely on the morphometric properties of the anterior commissure, a large and easily distinguishable white matter structure that is used as a reliable landmark for delineating the anterior hypothalamus border in other established protocols.

A second modification to existing protocols is the objective definition of the ventral border of the hypothalamus at the level of the infundibulum (pituitary stalk). This definition was easily identified and reliable, and allows for a more comprehensive study of the neuroendocrine axis, given that we included a definition of the ventral border of the stalk which allowed for reliable segmentation of the superior border of the pituitary gland. Although the stalk is a short and thin structure, which can be muddied by partial volume effects, the reliability of its segmentation was good, with both intra- and inter-rater Dice kappa overlap values of 0.79. The length and diameter of the stalk can be important clinical indications of infundibular disease (Satogami et al., 2010), and have been shown to increase across development (Sari et al., 2014), suggesting that inclusion of the pituitary stalk can be important in future structural MRI studies in developmental neuroendocrinology.

Third, the dorsal border of the hypothalamus within the tuberal region was slightly modified to include hypothalamic tissue medial to the fornix. This may prove to be a particularly important aspect of the protocol in study populations where the hypothalamic-pituitary-adrenal axis (i.e., the stress axis) is thought to play a role. The paraventricular nucleus (PVN) plays a key role in the stress response and energy balance.

Fourth, the proposed parcellation method divides the hypothalamus into periventricular, medial, and lateral zones. Although the hypothalamus is commonly divided into three compartments in the mediolateral (Saper et al., 2004; Schönknecht et al., 2013) or anteroposterior directions (Heiko Braak & Braak, 1992; Clark, 1936; Hofman & Swaab, 1989; Saper et al., 2004), existing manual segmentation protocols parcel the hypothalamus within its anteroposterior extent (Bocchetta et al., 2015; Makris et al., 2013). The current protocol proposes a mediolateral parcellation based on anatomical landmarks. As in the anteroposterior parcellations, the parcellation relies on geometric rules based on anatomical structures that are clearly identifiable on standard T1-weighted images, making the parcellation widely applicable. As such, this protocol extends the literature by providing a novel, detailed parcellation of the hypothalamus in the medial-lateral extent on high resolution structural MRIs.

As expected, sex differences in the total hypothalamic volume as well as within each hemisphere were detected. This sex difference is in line with others (Goldstein et al., 2007; Makris et al., 2013) and exemplifies the importance of further study of the hypothalamus in sexually differentiated health and disease states. Although Goldstein et al. (Goldstein et al., 2007) detected a larger right hypothalamus compared to the left, no hemispheric differences could be observed in our study. In line with other protocols however(Bocchetta et al., 2015; Makris et al., 2013; Stephanie Schindler et al., 2013), the right hypothalamus appeared slightly larger than the left in our sample.

While our regions of interest (the medial preoptic region, paraventricular region, and lateral hypothalamic region) were nominally larger in males compared to females, none of the regions were significant (the right preoptic region showed the strongest trend; Table 2, p=0.09). Outside our regions of interest, we found that the left posterior hypothalamic area was significantly larger in males than in females (Table 2, p=0.04), and the right anterior hypothalamus trended to be larger in males than in females (p=0.058). However, we did not detect a sex difference in the tuberal region, neither within the left or right hemisphere, nor in total volume (males: 530.3529, SD=120.41; females: 498.84, SD=119.71, p=0.44) which is in contrast to Makris et al. (Makris et al., 2013) who detected a sex difference in total volume. The effect size that we calculate from the Makris data is a Cohen’s d of 0.58, whereas our data estimate the effect size of the sex difference to be 0.26. We acknowledge however, that the Makris protocol may be more precise in the tuberal region, given that our protocol was focused on a parcellation method in the medial-lateral extent. True sex differences and effect size estimates will emerge as hypothalamic imaging, segmentation, and parcellation protocols become more detailed and refined as neuroimaging technology further advances.

Importantly, the sex differences in hypothalamic volume did not remain significant when covarying for salivary T levels. Although this finding suggests that T levels may account for the sex difference in hypothalamic volume, T was not correlated with hypothalamic volume. Thus, it is unclear what, other than the loss of power due to an additional covariate, led to the loss of significance in sex difference when covarying for T. Testosterone, acting via androgen receptors, is related to white matter growth across adolescence, with the effects being more pronounced in males than in females (J S Perrin et al., 2009; Jennifer S Perrin et al., 2008).

Because the hypothalamus is comprised of intermingled white and gray matter, it is possible that the sex difference in hypothalamic volume is primarily driven by white matter within it. Diffusion imaging has been applied within the hypothalamus (Schönknecht et al., 2013), and mean diffusivity appears to be higher in men than in women (Thomas et al., 2019). Further studies specifically aimed at determining sex differences in hypothalamic white matter structures and the dependency on testosterone and/or androgen receptors are needed to investigate this possibility.

On the other hand, the sex differences in hypothalamic volume remained when controlling for salivary estradiol. In fact, sex differences emerged, with men having a larger right MPA and posterior pituitary gland, but it is not clear what is driving this sex difference. A recent study reported that lower estradiol levels in girls aged 10-14 was associated with decreased gray matter volume, including within subcortical structures (i.e., the amygdala) across adolescence compared to girls with higher estradiol levels and this effect was not seen in boys (Herting et al., 2014). These data suggest that lower levels of estradiol in women can lead to decreased gray matter growth across adolescence. Most of the women in our study reported taking oral contraceptive pills, which leads to inhibitory feedback at the level of the hypothalamus and pituitary, preventing follicular growth and thus endogenous estradiol release. Thus, it is possible that prolonged oral contraceptive use blunted hypothalamic growth or reduced, which could account for the sex differences detected.

It is important to note that statistical methods to correct for total brain volume vary but regardless of the method, a sex differences is typically detected in hypothalamic volume. In our case, subject scans were analyzed in standardized space (ICBM 152) and therefore corrected for brain size. Some compute a ratio between the region of interest to total brain volume (Goldstein et al., 2001), others entering total brain volume as a covariate (Goldstein et al., 2001, 2002; Makris et al., 2013), and yet others correct by multiplying the region of interest volume by the inverse of the ratio of the group mean to the individual’s region of interest (Bocchetta et al., 2015). Thus, the sex difference in hypothalamic volume appears to be quite robust.

Our proposed hypothalamic segmentation protocol is most similar to, and complements that, of Bocchetta (Bocchetta et al., 2015). Our Dice kappa values were approximately 0.83 for left, right and whole hypothalamus, in comparison to Bocchetta (Bocchetta et al., 2015) who reported Dice values of 0.88 (SD=0.02) for the left and right hypothalamic hemispheres, using T1 and T2-weighted images acquired and segmented on scans with 1.1mm spatial resolutions. As such, despite using higher resolution images and extending hypothalamic boundaries, the present protocol is comparable in reproducibility to that of Bocchetta (Bocchetta et al., 2015). Mean Dice kappa intra-rater values for the POA, PVN and LH parcels ranged from 0.56-0.67, and we propose that the parcellation method described here can complement that originally defined by Makris et al. (Makris et al., 2013) and refined in Bocchetta (Bocchetta et al., 2015), depending on the region of interest. However, our protocol may be more appealing for studies interested in medial lateral divisions of the hypothalamus.

Although structural properties of the pituitary gland using MRI have been described developmentally (Castillo, 2005; MacMaster et al., 2007; Takano et al., 1999) and in psychopathological conditions (Delvecchio et al., 2017), there are some shortcomings to this work (as reviewed in (Anastassiadis et al., 2019)). First, the boundaries were previously not clearly defined, particularly at the level of the pituitary stalk, and between the anterior and posterior glands. In this protocol, we used the hypointense signal visible between the anterior and posterior pituitary glands, and found the posterior portion comprised an average of 24% of the total pituitary gland. This is consistent with histological reports that find the posterior gland comprised 20-30% of the entire pituitary gland (Garner et al., 2005), suggesting that the hypointense signal visible on T1-weighted images, and the hyperintense signal seen on T2- weighted images, may be a reliable and valid way to distinguish the two lobes.

Our proposed guidelines allow for a more detailed and analysis of the pituitary gland using high-resolution T1 and T2 weighted imaging, and provide a comprehensive approach to the morphological study of the hypothalamic-pituitary axes. As MRI technology advances to hone in on hypothalamic nuclei, a future direction could include analysis of different hypothalamic-pituitary-axes (e.g., the stress axis involving the PVN, and the reproductive axis involving the medial POA), its connections to other limbic structures and cortical regions (Lemaire et al., 2011; Saper et al., 2004; Schönknecht et al., 2013), and how those systems interact to induce structural changes in these systems that underlie various physiological conditions and behaviors.

This protocol can be applied in a variety of populations. The hypothalamus integrates somatic, visceral and behavioral information from the internal and external environment to coordinate autonomic responses and endocrine systems with the organisms behavior, including emotional responses (Noback et al., 2005). Nuclei such as the MPA, VMH, LH, and DMN are involved in reproductive, maternal, and feeding behaviors. The hypothalamus, and the preoptic region in particular, is also a key structure that may be involved in gender dysphoria (Garcia-Falgueras & Swaab, 2008; Hoekzema et al., 2015), yet has traditionally received little attention due to technical challenges of studying this structure in vivo. The hypothalamus is perhaps the most well-studied sexually dimorphic structure in the rodent brain, and is thus increasingly recognized as an important structure for understanding sex differences in psychopathological disorders, especially those that present sex/gender differences in onset, symptomatology and treatment, such as mood and anxiety disorders (Terlevic et al., 2013; Tognin et al., 2012), and psychosis (Goldstein et al., 2007; Tognin et al., 2012).

Future technological advances will surely lead to improved methods for the morphological study of the hypothalamus. As an integrative neural region, it has numerous afferents and efferents to and from its numerous specialized nuclei (Lemaire et al., 2011; Saper et al., 2004; Schönknecht et al., 2013), and as a result has intermingled white and gray matter, making its nuclei and subregions difficult to distinguish on 3T structural MRI images. The complex morphological and cytoarchitectural organization of the hypothalamus and its nuclei is also challenging using post-mortem histological preparations, as different histological stains reveal different nuclei and shapes of those nuclei (Bao & Swaab, 2011; Y Koutcherov et al., 2000; Yuri Koutcherov et al., 2007; Swaab & Fliers, 1985). These are further complicated by the numerous morphological sex and gender differences in volume and shape of certain nuclei (Hofman & Swaab, 1989; Swaab & Fliers, 1985). These technological challenges, due to the inherent complexity within the hypothalamus, might be at least partly addressed in the future, with higher in vivo resolution acquisitions, and as stronger MRI magnets becoming more readily accessible (Stephanie Schindler et al., 2013). Nonetheless, the advantage of a morphometric analysis of the hypothalamus using current parcellation methods on high resolution images acquired at 3T will improve our understanding of hypothalamic variations in vivo across time or under varying conditions.

We note that the parcellation method we applied to the mamillary bodies differs from manual segmentation protocols described by others (Copenhaver et al., 2006). A main difference is the exclusion of hyperintense voxels in our protocol within the hypothalamic region which likely represent major white matter tracts that connect to the mamillary bodies (i.e., fornix and mammillothalamic tract). This major methodological difference may make the parcellation of the mamillary bodies proposed in the current protocol less attractive to researchers, yet it may also represent a step forward in considering the white matter projections, as recently described for other limbic regions (Amaral et al., 2016).

This protocol and study have a number of strengths. First, the parcellation method complements and thus can be integrated into existing hypothalamic segmentation protocols proposed by others (Bocchetta et al., 2015; Makris et al., 2013). Second, it allows for a more comprehensive neuroendocrine analysis by adding the pituitary stalk and gland, as well as segmenting the functionally distinct anterior and posterior lobes, in addition to the posterior bright spot. A strength of the study is that we provide correlational analyses between two key sex steroid hormones and how they may be related to structural changes in the hypothalamus and pituitary.

A limitation of this study includes the small sample size. However, we had approximately equal numbers of men and women, and the effect size of the sex difference was similar to other reports. This suggests that the protocol can be reliably used to investigate sex differences. Nonetheless, the sample size may have limited our ability to detect sex differences in the parcels, associations between hypothalamic and pituitary regions, as well as with hormone levels, and to conduct sex specific analyses.

A second limitation is that the majority of women were taking oral contraceptives, limiting our ability to examine whether menstrual cycle phase and therefore fluctuating levels of endogenous estradiol influence hypothalamic/pituitary structures. Yet, we did collect salivary estradiol as well as testosterone, and we note that estradiol was correlated with brain volume as reported by others (Herting et al., 2014), but did not correlate with any hypothalamic measure. However, testosterone was correlated with the right PVN, suggesting that it can influence PVN volume when measured with MRI perhaps through actions at androgen receptors within the PVN (Clancy et al., 1992). Why these associations were detected in women but not men is unclear, but it would seem unlikely to be related to aromatase activity (which converts non-aromatizable androgens such as testosterone, into estradiol) given that no associations were detected with estradiol levels, and instead suggests a female-specific testosterone-dependent effect. Whether hypothalamic morphology is influenced by estradiol levels or exposure, as has been shown in the human hippocampus (Lord et al., 2008, 2010) as well as in the hypothalamus using animal studies (L H Calizo & Flanagan-Cato, 2000; Lyngine H Calizo & Flanagan-Cato, 2002; Ferri et al., 2014; Griffin & Flanagan-Cato, 2008), and whether it is affected by hormone administration (e.g., contraceptive pills and/or hormone replacement therapy following surgical, chemical, or natural menopause) warrants further study. Moreover, women should be sampled across the phases of the menstrual cycle, and in at least two consecutive cycles to increase validity of potential structural variations in hypothalamic volume.

In summary, we proposed a reliable, comprehensive segmentation protocol of the hypothalamus and pituitary gland, and detailed parcellations therein. Segmentation is done using T1 weighted images and accuracy is improved by acquisition of high-resolution T2- weighted images. We encourage researchers with both neuroimaging as well as neuroendocrine interests to employ this comprehensive approach to further the understanding of the role of the hypothalamus and pituitary in health and disease, while accounting for sex steroid hormones. Given that these structures are important in sexually differentiated endocrine systems and behaviors, applications of the protocol can also increase our understanding of sex and gender differences in health and diseases.

## Supporting information

Supplemental Materials Document 1 - HypPit_Thresholds_CleanDataSheet

## Acknowledgements

The authors gratefully acknowledge Suzanne King and David P. Laplante for access to the participants from their prospective longitudinal study, and supporting the implementation of the T2 neuroimaging acquisition used in the current segmentation protocol. We also thank Isabelle Bouchard for coordinating participant recruitment and scheduling, and Kyle McKee for assistance scanning the participants, as well as for technical assistance, including running BEaST via the BPIPE pipeline (available at https://github.com/CobraLab/minc-bpipe-library). We also thank Dorothee Schoemaker for help with image processing and useful discussions regarding the development of the protocol. We thank the lab of M. Mallar Chakravarty at the Douglas Research Center for technical support and assistance in the image acquisitions and processing, and access to their Computational Brain Imaging Lab’s open-source pipelines. Automated segmentation using BEaST was done on Compute Canada’s Advanced Research Computing Platforms, with technical assistance from Gabriel Devenyi. We also thank Alan Evans and his laboratory at the Montreal Neurological Institute, especially Paule-Joanne Toussaint, Claude Lepage, Chris Steele and Ayça Altinkaya for technical assistance viewing three dimensional histological BigBrain, and to Philippe Massicotte and Louis Borgeat at the National Research Council of Canada for support using the Atelier 3D software. We also thank Abbas Sadikot for discussions regarding anatomical boundaries within the hypothalamic region.

Funding was provided by a grant from the Canadian Institutes of Health Research (CIHR, FRN 125892) to Suzanne King, David P. Laplante, and Jens C. Pruessner. Sherri Lee Jones was funded by a postdoctoral fellowship provided by the Fonds de Québec – Santé (FRQ-S). The funding sources had no involvement in the collection, analysis or interpretation of the data, or writing of the report.

## Competing Interest

The authors have no competing interests to declare.

## Notes

### Competing Interest Statement

The authors have declared no competing interest.

