## Supplemental Materials Document 1 - HypPit_Thresholds_CleanDataSheet for "Effects of sex and gonadal hormones on manually segmented hypothalamic and pituitary gland volumes in young healthy adults"

| | HYPOTHALAMUS Contrast T1 (T2) | | HYPOTHALAMUS Contrast (T2) | | PITUITARY Contrast T1 (T2) | | Hypothalamus Gray Matter min threshold 40% of range | Hypothalamus Gray Matter max threshold (75% of range) | Pituitary | | | |
|---|---|---|---|---|---|---|---|---|---|---|---|---|
| ID | Min | Max | Min | Max | Min | Max | | | Gray matter min | Bright Spot (75%) | Alignment of T1/T2 | Notes |
| | | | | | | | 0 | 0 | 0 | 0 | | |
| | | | | | | | 0 | 0 | 0 | 0 | | |
| | | | | | | | 0 | 0 | 0 | 0 | | |
| | | | | | | | 0 | 0 | 0 | 0 | | |
| | | | | | | | 0 | 0 | 0 | 0 | | |
| | | | | | | | 0 | 0 | 0 | 0 | | |
| | | | | | | | 0 | 0 | 0 | 0 | | |
| | | | | | | | 0 | 0 | 0 | 0 | | |
| | | | | | | | 0 | 0 | 0 | 0 | | |
| | | | | | | | 0 | 0 | 0 | 0 | | |
| | | | | | | | 0 | 0 | 0 | 0 | | |
| | | | | | | | 0 | 0 | 0 | 0 | | |
| | | | | | | | 0 | 0 | 0 | 0 | | |
| | | | | | | | 0 | 0 | 0 | 0 | | |
| | | | | | | | 0 | 0 | 0 | 0 | | |
| | | | | | | | 0 | 0 | 0 | 0 | | |
| | | | | | | | 0 | 0 | 0 | 0 | | |
| | | | | | | | 0 | 0 | 0 | 0 | | |
| | | | | | | | 0 | 0 | 0 | 0 | | |
| | | | | | | | 0 | 0 | 0 | 0 | | |
| | | | | | | | 0 | 0 | 0 | 0 | | |
| | | | | | | | 0 | 0 | 0 | 0 | | |
| | | | | | | | 0 | 0 | 0 | 0 | | |
| | | | | | | | 0 | 0 | 0 | 0 | | |
| | | | | | | | 0 | 0 | 0 | 0 | | |
| | | | | | | | 0 | 0 | 0 | 0 | | |
| | | | | | | | 0 | 0 | 0 | 0 | | |
| | | | | | | | 0 | 0 | 0 | 0 | | |
| | | | | | | | 0 | 0 | 0 | 0 | | |
| | | | | | | | 0 | 0 | 0 | 0 | | |
| | | | | | | | 0 | 0 | 0 | 0 | | |
| | | | | | | | 0 | 0 | 0 | 0 | | |
| | | | | | | | 0 | 0 | 0 | 0 | | |
| | | | | | | | 0 | 0 | 0 | 0 | | |
| | | | | | | | 0 | 0 | 0 | 0 | | |
| | | | | | | | 0 | 0 | 0 | 0 | | |
| | | | | | | | 0 | 0 | 0 | 0 | | |
| | | | | | | | 0 | 0 | 0 | 0 | | |
| | | | | | | | 0 | 0 | 0 | 0 | | |
| | | | | | | | 0 | 0 | 0 | 0 | | |
| | | | | | | | 0 | 0 | 0 | 0 | | |
| | | | | | | | 0 | 0 | 0 | 0 | | |
| | | | | | | | 0 | 0 | 0 | 0 | | |
| | | | | | | | 0 | 0 | 0 | 0 | | |
| | | | | | | | 0 | 0 | 0 | 0 | | |
| | | | | | | | 0 | 0 | 0 | 0 | | |
| | | | | | | | 0 | 0 | 0 | 0 | | |
| | | | | | | | 0 | 0 | 0 | 0 | | |
| | | | | | | | 0 | 0 | 0 | 0 | | |
| | | | | | | | 0 | 0 | 0 | 0 | | |
| | | | | | | | 0 | 0 | 0 | 0 | | |
| | | | | | | | 0 | 0 | 0 | 0 | | |
| | | | | | | | 0 | 0 | 0 | 0 | | |
| | | | | | | | 0 | 0 | 0 | 0 | | |
| | | | | | | | 0 | 0 | 0 | 0 | | |
| | | | | | | | 0 | 0 | 0 | 0 | | |
| | | | | | | | 0 | 0 | 0 | 0 | | |
| | | | | | | | 0 | 0 | 0 | 0 | | |
| | | | | | | | 0 | 0 | 0 | 0 | | |
| | | | | | | | 0 | 0 | 0 | 0 | | |
| | | | | | | | 0 | 0 | 0 | 0 | | |
| | | | | | | | 0 | 0 | 0 | 0 | | |
| | | | | | | | 0 | 0 | 0 | 0 | | |
| | | | | | | | 0 | 0 | 0 | 0 | | |
| | | | | | | | 0 | 0 | 0 | 0 | | |
| | | | | | | | 0 | 0 | 0 | 0 | | |
| | | | | | | | 0 | 0 | 0 | 0 | | |
| | | | | | | | 0 | 0 | 0 | 0 | | |
| | | | | | | | 0 | 0 | 0 | 0 | | |
| | | | | | | | 0 | 0 | 0 | 0 | | |
| | | | | | | | 0 | 0 | 0 | 0 | | |
| | | | | | | | 0 | 0 | 0 | 0 | | |
| | | | | | | | 0 | 0 | 0 | 0 | | |
| | | | | | | | 0 | 0 | 0 | 0 | | |
| | | | | | | | 0 | 0 | 0 | 0 | | |
| | | | | | | | 0 | 0 | 0 | 0 | | |
| | | | | | | | 0 | 0 | 0 | 0 | | |
| | | | | | | | 0 | 0 | 0 | 0 | | |
| | | | | | | | 0 | 0 | 0 | 0 | | |
| | | | | | | | 0 | 0 | 0 | 0 | | |
| | | | | | | | 0 | 0 | 0 | 0 | | |
| | | | | | | | 0 | 0 | 0 | 0 | | |
| | | | | | | | 0 | 0 | 0 | 0 | | |
| | | | | | | | 0 | 0 | 0 | 0 | | |
| | | | | | | | 0 | 0 | 0 | 0 | | |
| | | | | | | | 0 | 0 | 0 | 0 | | |
| | | | | | | | 0 | 0 | 0 | 0 | | |
| | | | | | | | 0 | 0 | 0 | 0 | | |
| | | | | | | | 0 | 0 | 0 | 0 | | |
| | | | | | | | 0 | 0 | 0 | 0 | | |
| | | | | | | | 0 | 0 | 0 | 0 | | |
| | | | | | | | 0 | 0 | 0 | 0 | | |
| | | | | | | | 0 | 0 | 0 | 0 | | |
| | | | | | | | 0 | 0 | 0 | 0 | | |
| | | | | | | | 0 | 0 | 0 | 0 | | |
| | | | | | | | 0 | 0 | 0 | 0 | | |
| | | | | | | | 0 | 0 | 0 | 0 | | |
| | | | | | | | 0 | 0 | 0 | 0 | | |
| | | | | | | | 0 | 0 | 0 | 0 | | |
| | | | | | | | 0 | 0 | 0 | 0 | | |
| | | | | | | | 0 | 0 | 0 | 0 | | |
| | | | | | | | 0 | 0 | 0 | 0 | | |
| | | | | | | | 0 | 0 | 0 | 0 | | |
| | | | | | | | 0 | 0 | 0 | 0 | | |
| | | | | | | | 0 | 0 | 0 | 0 | | |
| | | | | | | | 0 | 0 | 0 | 0 | | |
| | | | | | | | 0 | 0 | 0 | 0 | | |

To continue beyond this point, select the four cells above, and apply the formulas (drag) to next cells
